# Sleep neural code perpetuates the evolving negativity bias under stress

**DOI:** 10.64898/2026.03.03.709197

**Authors:** Seung Min Um, Reiko Okubo-Suzuki, Masanori Nomoto, Mahmoud Khaled Hanafy, Murayama Emi, Kaori Yamada-Nomoto, Hirotaka Asai, Kiriko Choko, Akinobu Suzuki, Kaoru Inokuchi

**Affiliations:** Department of Biochemistry, Graduate School of Medicine and Pharmaceutical Sciences, University of Toyama, Toyama, 930−0194, Japan; Research Centre for Idling Brain Science, University of Toyama, Toyama, 930−0194, Japan; Department of Anatomy and Cell Biology, Schulich School of Medicine and Dentistry, Western University, London, Ontario N6A 3K7, Canada; Graduate School of Medicine, The University of Tokyo, Tokyo, Japan; Learning and Memory Project, Tokyo Metropolitan Institute of Medical Science, Tokyo 156-8506, Japan

## Abstract

Negativity bias is the distorted cognitive processing of how self-experiences are negatively perceived through rumination and overgeneralization. Despite proposed theories of higher-order emotional representation and the involvement of default mode networks, the biological mechanisms underlying how negative emotional bias is persistently generated remain unknown. Here, we show that under negative emotional state, sleep orchestrates neural dynamics to shape persistent negativity bias. A negative emotional state, induced by repeated social defeat stress (SoD), generates negativity bias in both memory–recall and novel experiences such as fear conditioning. Longitudinal Ca^2+^ imaging of the hippocampus across days revealed that in emotional states, negative schema-like representations that encode the semantic aspect of negativity across experiences are generated. These negative schema-like representations are continuously processed along with dynamic co-reactivations of distinct emotional experiences during sleep. Closed-loop optogenetic silencing of stress experience-tagged cellular ensembles during sleep, but not during awake, prevents negativity bias in future memory–recall and novel experiences. Finally, sleep-active, but not sleep-nonactive, SoD cells predict and decode emotional behaviours during learned and novel situations. Together, these results reveal the biological emergence of semantic-like representations of emotional experiences and sleep causally coordinate neuronal dynamics to persistently shape negativity in future conscious states. These findings offer a new perspective on how higher-order emotional processing during sleep may determine our emotional interpretation of past and future experiences.

## Introduction

Negativity bias, a core feature of mood disorders such as anxiety and depression, distorts perceptions of self and the world towards pessimism^1,2^. This cognitive distortion drives a vicious cycle of negative self-reflection, rumination, and overgeneralized autobiographical experiences, which chronically worsens mental well-being^3^. Influential psychological models link subjective emotional interpretation to cognitive brain functions. Beck, who developed the Beck Depression Inventory and pioneered cognitive behavioural therapy (CBT), proposed that depression stems from internal negative schemas, i.e., negative mental representations interplaying with biased attention and memory in higher-order areas^4^. One model suggests that these cognitive regions encode conscious emotional experiences, resulting in the generation of subjective feelings such as fear^5^.

Recent neuroimaging studies have revealed that resting-state connectivity and default mode networks, which consist of/interact with higher-order and memory regions^6^, are associated with perpetual negative emotional processing, such as generalized anxiety disorder^7^, self-referential processing^8^ and rumination^9,10^. Rodent studies have shown that sharp-wave ripples during rest modulate stress-impaired sociability^11^, that rapid-eye-movement (REM) sleep activity regulates fear memory^12^, and that midbrain circuit induces stress-driven restorative sleep^13^. These findings emphasize the conserved role of offline states in emotional defensive regulation.

However, how cognitive representations of emotional states, such as schema-like constructs reflecting the concept of negativity evolved by experiences, are generated and sustained remains elusive. Previous studies have indicated that the cognitive brain area consolidates memories offline^14–16^ and integrates experiences into gist-like structures or preexisting knowledge for adaptive cognition^17–20^. We hypothesize that cognitive memory networks create neuronal representations of semantic negativity from autobiographical experiences, with offline states enabling updates that are potentially biased towards overgeneralization in intense negative states. We targeted the dorsal hippocampus, a region critical for both memory and higher-order cognition^21,22^, which processes multimodal cognitive maps of episodic experiences (spatial and nonspatial)^23–25^ and forms synchronous networks with cortical areas during offline states^26–28^, a function that is conserved across species.

## Results

### Induction of Negative Emotional States Biases Interpretations of Congruent and Novel Experiences

We developed a rodent behavioural paradigm mimicking emotional state-dependent negativity bias in humans. In the task, a stable negative emotional state is first induced through repeated social defeat stress (SoD), which is known to recapitulate the core features of mood disorders such as social avoidance and anxiety^29–32^. The SoD mice encountered aggressive retired breeder ICR mice for 2–3 min daily, whereas the neutral group explored a novel home cage. We then conducted a social interaction (SI) test in an open field arena, with a novel ICR mouse absent or present in a wire-mesh cage (Fig. 1a). Compared with the neutral mice, the SoD mice spent less time in the interaction zone and more time in opposite corners, indicating social avoidance behaviour (Fig. 1b–c and Extended Data Fig. 1a–b). Baseline anxiety was assessed via the elevated plus maze (EPM) test one day post-SoD (Fig. 1d). Compared with the mice in the neutral group, the mice in the SoD group explored the open arms less frequently, confirming the enhanced anxiety-like behaviour of the mice in this group (Fig. 1e–f). Therefore, a six-day SoD paradigm induces a stable negative emotional state beyond the transient arousal effect.

**Fig. 1.**
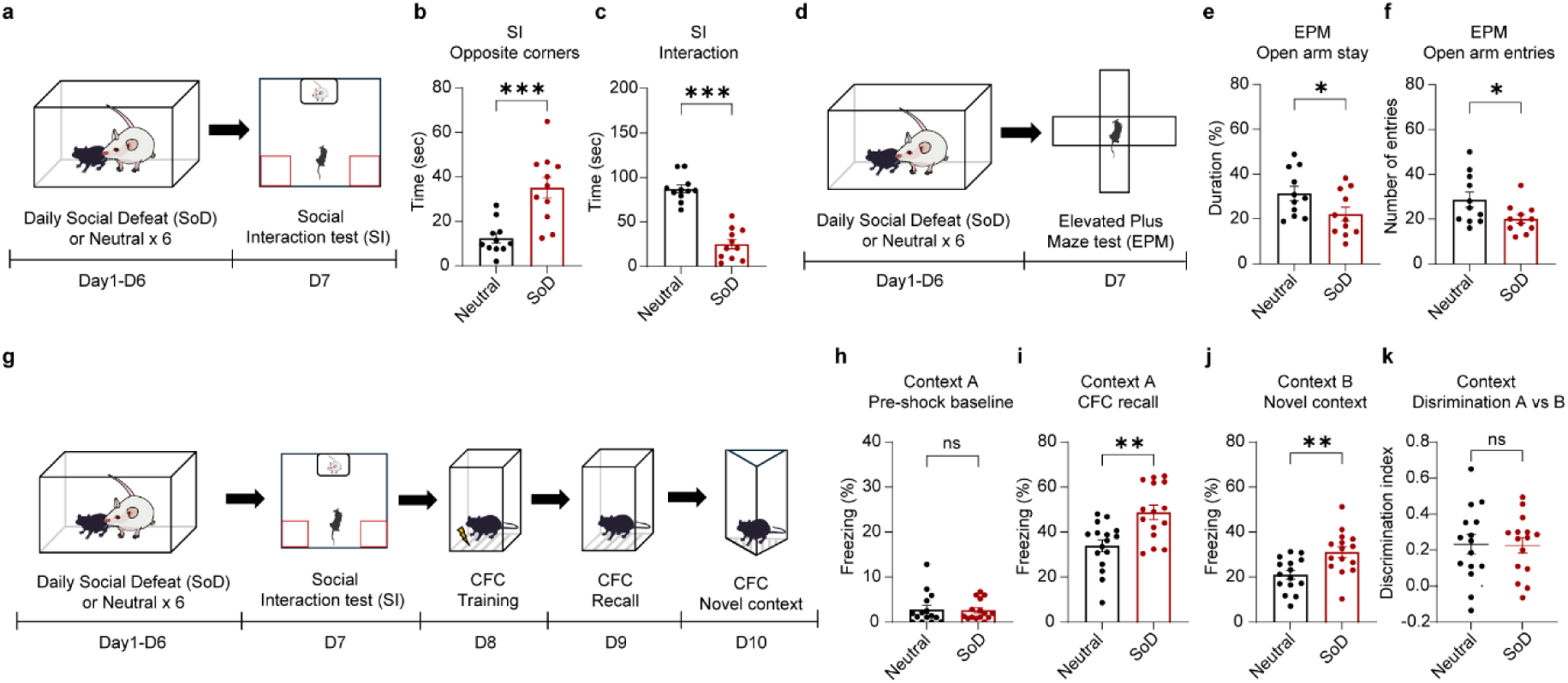
Repeated social defeat stress (SoD) drives social avoidance, anxiety, and emotional bias in future memory recall and novel experiences. **a.** Schematic of the behavioural experiments showing repeated social defeat stress (SoD) followed by the social interaction (SI) test. **b.** Time spent in opposite corners during novel ICR mouse encounters during the SI test (SI-ICR) (Neutral *n* = 11; SoD *n* = 11 mice; unpaired t test with correction, *t* = 4.374; *P* < 0.001). **c**. Time spent in the interaction zone during the SI-ICR session (Unpaired t test, *t* = 9.029, *P* < 0.001). **d.** Schematic of the timeline of the SoD phase followed by the elevated plus-maze test (EPM) for testing anxiety. **e.** Percentage of time spent in the open arms during the EPM test (Neutral *n* = 11; SoD *n* = 11 mice; unpaired t test, *t* = 2.093; *P* = 0.0493). **f.** Total number of entries into the open arm in the EPM test (Unpaired t test, *t* = 2.187, *P* = 0.0408). **g**. Timeline of the behavioural schedule for the SoD-SI test followed by a contextual fear conditioning (CFC) memory test and novel context exploration. **h.** Percentage of total freezing time during pre-shock exploration for 2 minutes (Neutral *n* = 15; SoD *n* = 15 mice; Mann‒Whitney *U* test, *P* = 0.6891). **i.** Percentage of total freezing time during the CFC memory recall test (Context A) (Unpaired t test, *t* = 3.379, *P* = 0.0022). **j.** Percentage of total freezing time during novel context exposure (Context B) (Unpaired t test, *t* = 3.229, *P* = 0.0032). **k.** Discrimination index calculated according to the freezing levels measured in contexts A and B (Unpaired t test, *t* = 0.0956, *P* = 0.9246). Data are presented as the mean ± s.e.m. Statistical significance is expressed as follows: ns, not significant; * *P* < 0.05; ** *P* < 0.01; *** *P* < 0.001.

We then examined how this negative state influences the interpretation of new emotional experiences. The mice were subjected to contextual fear conditioning (CFC) one day post-SI; the paradigm involved 2 min of square chamber exploration followed by electric shock, and fear memory was tested 24 h later (Fig. 1g). Baseline freezing during pre-shock exploration did not differ between the SoD group and the neutral group (Fig. 1h). However, during the recall phase, compared with the neural mice, the SoD mice exhibited substantially greater freezing behaviour, indicating an enhanced emotional response (Fig. 1i). A short-term memory (STM) test was conducted 30 min after training, and no group differences were identified, highlighting the need for a post-training incubation period (Extended Data Fig. 1c–d). When mice explored a novel context with altered wall patterns but similar grid floors, SoD mice showed elevated freezing behaviour in the novel chamber relative to neutral mice (Fig. 1j), suggesting negatively biased perception across learned and novel environments. Comparable freezing reductions from learned to novel contexts and similar discrimination indices ruled out impaired context differentiation as the cause (Fig. 1k and Extended Data Fig. 1e–f).

To test the specificity of the emotional state, we induced a positive reward state, female social interaction^33,34^ (FSI), for six days in male mice, followed by the CFC protocol (Extended Data Fig. 1g). In contrast to the increased freezing in the SoD group, the FSI group displayed reduced freezing during recall compared with that of the neutral group (Extended Data Fig. 1h). Together, these results suggest that the repeated SoD stress protocol sufficiently induced a negative emotional state in rodents, mimicking human-like negativity, and that this state biases the interpretation of emotionally congruent experiences.

### Emergence of Schema-Like Neuronal Representations Encoding Generalized Negativity

We next investigated how neural representations of negativity bias evolve through emotional state induction and engage with congruent experiences across days. On the basis of the role of higher-order brain functions in memory networks that build complex multimodal cognitive maps with semantic and episodic components^35^, we hypothesized that memory-related areas generate specific representations encoding the subjective concept of emotion shaped by episodic experiences (Fig. 2a). We injected the Janelia variant Ca^2+^ indicator jGCaMP7b into the dorsal hippocampal CA1 region of the mice and conducted longitudinal Ca^2+^ imaging across days, thereby enabling the tracking of stable neural representations across multiple emotional experiences (Fig. 2b–c).

**Fig. 2.**
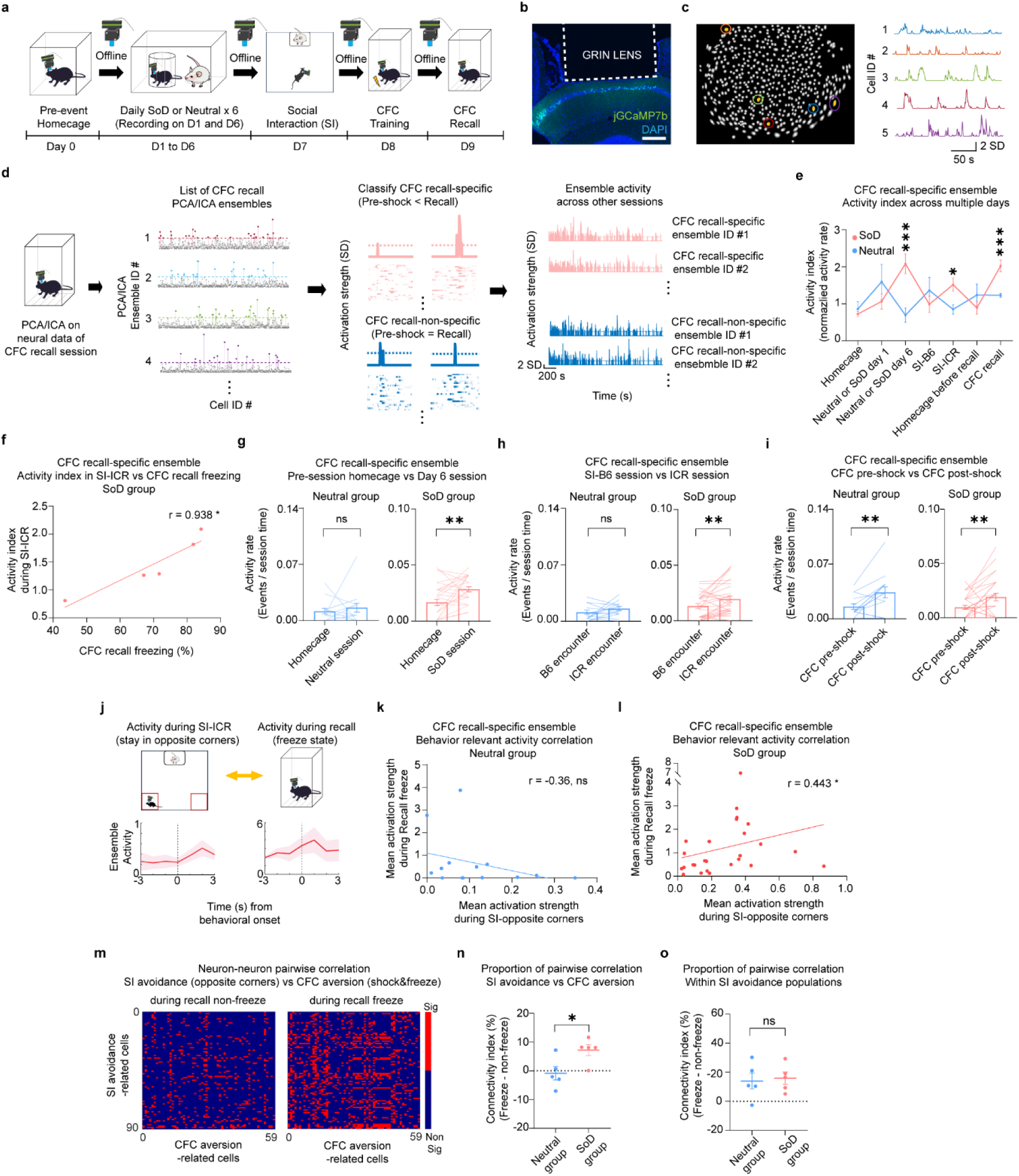
Longitudinal Ca^2+^ imaging analysis showing the emergence of negative schema-like ensembles and increased functional connectivity among subtypes of emotional behaviour-related cells in the SoD mice. **a.** Schematic of longitudinal Ca^2+^ imaging recording schedules from the SoD experience session to the CFC memory recall session. **b.** Expression of the Janelia variant Ca^2+^ indicator AAV9-hSyn-JGCaMP7b and the location of the implanted GRIN lens in the CA1 region of the dorsal hippocampus. Scale bar, 250 µm. **c.** Example image of cells extracted from Ca^2+^ imaging and representative traces of z scored neural activity. **d.** Schematic of the analysis flowchart for detecting recall-specific ensembles using PCA/ICA dimensionality reduction methods. After all PCA/ICA ensembles from the CFC recall session were detected, these ensembles were further categorized into recall-specific (pink) or recall-nonspecific ensembles (blue) by comparing the ensemble activity in the CFC recall session with that in the previous CFC pre-shock session by Wilcoxon rank sum methods. For each example trace of the ensemble activity, the corresponding raster plot of the neural activity is shown with the same colour. **e.** Multisession comparison of the averaged activity index of the recall-specific ensembles, which is normalized by the average activity rate of the recall-nonspecific ensembles per mouse for each corresponding session. (Neutral *n* = 14; SoD *n* = 28 ensembles; two-way RM ANOVA with correction; interaction *P* <0.001; *F*(6, 240) = 5.382; Šídák’s multiple comparisons test). **f.** Pearson correlation analysis of the activity index of recall-specific ensemble during the SI-ICR session and prospective total freezing level (%) in the CFC recall session per SoD mouse. (*r* = 0.9378, *P* = 0.0184, *n* = 5 mice) **g.** Activity rate of each CFC recall-specific ensemble in the neutral group in the home cage before the neutral experience (Pre-Neutral-home cage) and during the neutral session (novel home cage exploration) on day 6 (*n* = 14 ensembles; Wilcoxon signed rank test; *P* = 0.3306); additionally, the activity rate of the SoD group in the home cage before the SoD session (Pre-SoD-home cage) and during the SoD session on day 6 of the SoD experience (*n* = 28 ensembles, Wilcoxon signed rank test, *P* = 0.0014). **h.** Activity rate of each recall-specific ensemble during the novel B6 mouse encounter (SI-B6) and novel ICR mouse encounter (SI-ICR) in the social interaction test for the neutral group (*n* = 14 ensembles, Wilcoxon signed rank test, *P* = 0.4352) and SoD group (*n* = 28 ensembles, Wilcoxon signed rank test, *P* = 0.0066). **i.** Activity rate of each recall-specific ensemble during the 2-minute exploration period before receiving the first electric shock (CFC pre-shock) and during the remaining session after the delivery of the first shock (CFC post-shock) on the CFC conditioning day in the neutral group (*n* = 14 ensembles; Wilcoxon signed rank test; *P* = 0.0076) and the SoD group (*n* = 28 ensembles, Wilcoxon signed rank test, *P* = 0.0053). **j.** Schematic for assessing the correlation of the CFC recall-specific ensemble activity during opposite corner stays during the SI-ICR session and the ensemble activity during the freeze state in the CFC recall session. Example traces of averaged recall-specific ensemble activity (z scored) around the onset of the opposite corner entries in the SI-ICR (lower left) and during freezing behaviours in CFC recall (lower right) sessions. **k.** Spearman correlation (*r* value shown) for the activity of the CFC recall-specific ensembles in the neutral group during opposite corner stays and activity during freezing in the CFC recall session for the neutral group (*r* = - 0.3600, *P* = 0.2047) and **l.** SoD group (*r* = 0.4426, *P* = 0.0208, *n* = 27 ensembles). **m.** Representative heatmaps showing the intercell type pairwise correlation analysis between SI avoidance-related (SI opposite corner-related neurons) and CFC aversion-related (shock- and freeze-related neurons) cells in the non-freezing state (left) and the freezing state in the CFC recall session (right). Red indicates a significant correlation between two neurons. **n.** Percentage of significant pairwise correlations of SI avoidance-related and CFC aversion-related cells during the freezing state in the CFC recall session (Neutral *n* = 5; SoD *n* = 5 mice; unpaired t test, t = 2.653; *P* = 0.0291). **o.** Percentage of significant intracell type neuron‒neuron pairwise correlations of SI avoidance-related neurons during the freezing state in the CFC recall session (Neutral, *n* = 5; SoD, *n* = 5 mice; Unpaired t test, *t* = 0.2808; *P* = 0.786). Data are presented as the mean ± s.e.m. Statistical significance is expressed as * *P* < 0.05; ** *P* < 0.01; and *** *P* < 0.001.

To extract neural representations encoding generalized negativity across episodic experiences, we targeted neural activity during the CFC recall session, where SoD mice exhibit increased freezing due to the prior SoD paradigm. PCA/ICA dimensionality reduction was used to identify cell ensembles that were active during the CFC recall session (Fig. 2d and Extended Data Fig. 2a–b). We defined CFC recall-specific ensembles as those showing significantly higher activity in the CFC recall session (CFC Recall) than in the CFC habituation session before the first electric shock (CFC Pre-shock). There was no difference in the total number of CFC recall ensembles between the neutral and SoD groups (Extended Data Fig. 2c), whereas the proportion of CFC recall-specific ensembles shows increased tendency in the SoD group despite the lack of significant difference (Extended Data Fig. 2d).

When the activity index of the CFC recall-specific ensembles (normalized by the mean activity rate of the CFC recall-nonspecific ensembles) was compared with the neutral mice across the days, the SoD mice displayed an increased ensemble activity index only during negative emotional experiences, such as on the last day of social defeat (SoD day 6) and when they encountered stranger ICR mice during social interactions (SI-ICR) (Fig. 2e). This suggests the emergence of emotional schema-like neuronal representations in the SoD group, where ensemble activity is likely driven by the concept of negativity rather than specific contexts of single experiences. Indeed, ensemble activity index in the SI-ICR session were positively correlated with the total freezing levels in the CFC recall sessions for individual SoD mice, implying that avoidance-related activity predicts future negativity bias in new experiences (Fig. 2f). Within-group session-to-session analysis of the activity rates of the CFC recall-specific ensembles revealed that compared with the pre-SoD home cage sessions, the SoD group displayed increased activation during the SoD sessions, while the neutral group exhibited no observable difference in activity rates between the novel home cage exploration and pre-exploration home cage sessions (Fig. 2g). We next compared the ensemble activity rates between sessions in which the mice encountered stranger B6 mice (SI-B6) and stranger ICR mice (SI-ICR) in wire-mesh cages. Compared with that in SI-B6 session, the SoD group showed increased CFC recall-specific ensemble activation in SI-ICR session, whereas no observable change in activation was observed in the neutral group (Fig. 2h), suggesting that the CFC recall-specific ensembles in the SoD group reflect negative emotional states (social avoidance) rather than general social stimuli. Compared with that in the pre-shock session, the activity of the CFC recall-specific ensemble in both groups increased during the post-shock session (Fig. 2i), indicating sensitivity to aversive inputs in the dorsal hippocampus.

Significant correlations between ensemble activities emerged during opposite-corner stays, which are associated with social avoidance, and during freezing in the CFC recall session in the SoD group but not in the neutral group (Fig. 2j–l). This result strongly suggests that CFC recall-specific ensembles encode negative schema-like information, i.e., generalized negativity, independent of behavioural subtypes and environmental specificity.

To investigate interactions among subsets of emotional behaviour-related cells (Extended Data Fig. 3a–c) during the biased response in the CFC recall session, we computed intercell type pairwise correlations between SI avoidance-related (opposite-corner stay) cells and CFC aversion-related (shock and freeze) cells during freezing in the CFC recall session (Fig. 2m). The proportion of significantly correlated pairs was greater in the SoD group than in the neutral group (Fig. 2n), with no group difference in significant intracell type pairwise correlations among SI avoidance-related cells (Fig. 2o), suggesting increased coordination between social avoidance-related and fear-related populations during freezing in the CFC recall session. Interestingly, the overlapping proportions of SI avoidance-related cells and CFC aversion-related cells were comparable between the groups (Extended Data Fig. 3d). Together, these results reveal that negative emotional states lead to the emergence of schema-like representations encoding semantic aspects of negativity, with coordinated interactions among different subtypes of negative emotion-related cells during biased behavioural responses. This suggests that cognitive memory networks form representations of experiential negativity in an emotional state-dependent manner.

### Sleep Activates Schema-Like Ensembles and Drives the Interaction of Congruent Emotional Experiences

We next investigated how schema-like representations of negativity are maintained and propagated to future emotional states, driving negative overgeneralization. Given that rumination and interpretation biases are repetitive negative processes that occur in the off-task resting state^1,9^, we hypothesized that neural subpopulations representing generalized negativity are continuously processed and engaged after new emotionally similar events. We quantified the activity rate during offline periods of the CFC recall-specific ensembles with EEG and EMG recordings (Fig. 3a–b). Compared with that in the pre-neutral awake state, the ensemble activity rate in the post-CFC-training awake state (30 min after home cage return) increased in the neutral group (Fig. 3c), whereas no significant difference was observed in the SoD group (Fig. 3d). With respect to NREM sleep comparisons, activity rates increased significantly from the post-SI NREM (SI-NREM) state to the post-CFC NREM (CFC-NREM) state in both groups, but no notable change was observed from the pre-neutral/pre-SoD NREM state to the post-CFC NREM state (Fig. 3e–f). Notably, during REM sleep, the activity rate of the SoD group robustly increased in the post-CFC REM state compared with that in the pre-SoD REM and post-SI REM states, but this trend was absent in the neutral group (Fig. 3g–h). These findings indicate that a negative emotional state in SoD mice drives the continuous processing of schema-like representations during offline periods, specifically in REM sleep.

**Fig. 3.**
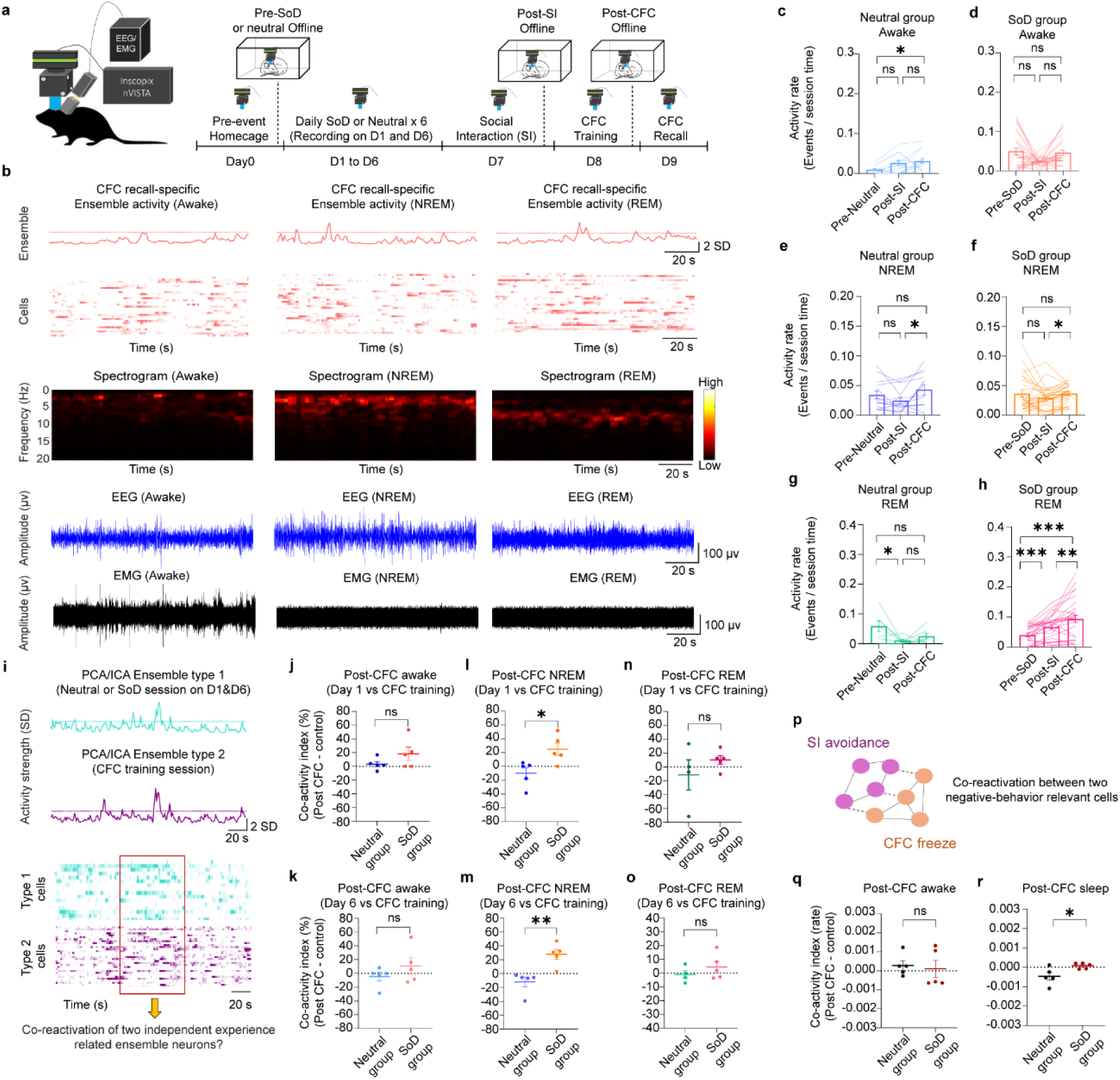
Offline sleep activates and coordinates negativity representations. **a.** Schematic of offline recording sessions during the SoD (or neutral) CFC recall experimental paradigm. **b.** Examples of traces of CFC recall-specific ensemble activity during offline awake, offline NREM sleep and REM sleep periods with an ensemble neuron raster map, representative heatmaps of spectrograms and activity traces of EEG and EMG recordings for the corresponding offline sessions. **c.** CFC recall-specific ensemble activity rate of the neutral group during offline awake sessions from the control home cage session before the neutral experience (Pre-Neutral) and the home cage session after the SI (post-SI) and CFC conditioning (post-CFC) sessions (*n* = 14 ensembles; Friedman test, *P* = 0.0095; Dunn’s multiple comparisons test). **d.** CFC recall-specific ensemble activity rate of the SoD group during offline awake sessions, including the home cage before the SoD experience, the post-SI period and the post-CFC period (n = 28 ensembles; Friedman test, *P* = 0.0649; Dunn’s multiple comparisons test). **e.** CFC recall-specific ensemble activity rate of the neutral group in NREM sessions (*n* = 14 ensembles, Friedman test, *P* = 0.0244, Dunn’s multiple comparisons test) and **f.** CFC recall-specific ensemble activity rate of the SoD group in NREM sessions (*n* = 28 ensembles; Friedman test, *P* = 0.0455; Dunn’s multiple comparisons test). **g.** CFC recall-specific ensemble activity rate of the neutral group in REM sessions (*n* = 7 ensembles; one-way RM ANOVA; *P* = 0.0194; *F* (2, 12) = 5.574; Tukey’s multiple comparisons test), and **h.** CFC recall-specific ensemble activity rate of the SoD group in REM sessions (*n* = 28 ensembles, one-way RM ANOVA with correction; *P* < 0.001; *F* (1.355, 36.58) = 20.70; Tukey’s multiple comparisons test). **i.** Schematic of the co-reactivation of two emotional experiences: one from SoD (or neutral) session-related PCA/ICA ensemble neurons and another from CFC training session-related PCA/ICA ensemble neurons. **j**. Co-reactivation of SoD (or neutral) day 1 and CFC-training ensemble neurons in the post-CFC awake state (Neutral *n* = 5; SoD *n* = 5 mice; unpaired t test, *t* = 1.432; *P* = 0.1901). **k**. Co-reactivation of SoD (or neutral) day 6 and CFC-training ensemble neurons in the post-CFC awake state (Unpaired t test, *t* = 1.159, *P* = 0.2797). **l**. Co-reactivation of SoD (or neutral) day 1 (unpaired t test, t = 2.852, *P* = 0.0214), **m.** SoD (or neutral) day 6 and CFC-training ensemble neurons in the post-CFC NREM sleep state (Mann–Whitney U test, *P* = 0.0079). **n**. Co-reactivation of SoD (or neutral) day 1 (Neutral *n* = 4; SoD *n* = 5 mice; unpaired t test, t = 1.051; *P* = 0.3283), **o.** SoD (or neutral) day 6 and CFC-training ensemble neurons in the post-CFC REM sleep state (unpaired t test, *t* = 1.021; *P* = 0.3413). **p.** Schematic of the analysis of the offline co-reactivation of two emotional behaviour-related cells: SI avoidance-related and CFC freeze-related neurons. **q.** Co-reactivation of SI avoidance-related and CFC freeze-related neurons in the post-CFC awake state (Neutral *n* = 5; SoD *n* = 5 mice; Mann–Whitney *U* test; *P* = 0.5476). **r.** Co-reactivation of SI avoidance-related and CFC freeze-related neurons in the post-CFC sleep state (unpaired t test; *t* = 2.567; *P* = 0.0333). Data are presented as the mean ± s.e.m. Statistical significance is expressed as follows: ns, not significant; * *P* < 0.05; ** *P* < 0.01; *** *P* < 0.001.

Building on these results, we explored dynamic interactions between past and new episodic experience-related populations, which are contextually distinct and temporally separated but share only emotional commonality. We applied PCA/ICA independently to the neural data of the mice from neutral or SoD sessions on day 1 or day 6, as well as to the data from the CFC training session, to extract ensembles for past (neutral/SoD) and new (CFC) episodic experiences (Fig. 3i). To quantify how these ensembles interact during offline periods, we computed the proportion of significant pairwise co-reactivations (co-activity index) between these ensemble cells during post-CFC offline periods, normalized by subtracting pre-experience (neutral or SoD) offline proportions (Extended Data Fig. 4a). Consistent with the CFC recall-specific ensemble dynamics in the prior offline-awake analyses (Fig. 3c–d), no notable differences in the co-reactivation of neutral or SoD session-related ensemble cells (on day 1 or day 6) and CFC training session-related ensemble cells in the post-CFC awake state (Fig. 3j–k) were observed in either group.

When the co-reactivation analysis was extended to sleep sessions, however, compared with the neutral group, the SoD group showed substantial increases in co-reactivation during post-CFC NREM sleep between SoD session-related ensemble cells (either day 1 or 6) and CFC training session-related ensemble cells (Fig. 3l–m). No significant co-reactivations appeared during REM sleep (Fig. 3n-o), in contrast to the dynamics of the schema-like recall-specific ensembles during REM sleep (Fig. 3h). These results suggest that emotionally similar, yet distinct, episodic cell populations robustly interact during NREM sleep, highlighting the role of NREM sleep in guiding dynamic interactions across broad emotional experiences. Interestingly, we observed an increased proportion of pairwise correlations at the single-cell level between co-reactivated SoD ensemble cells (which co-reactivated with CFC training ensemble cells during post-CFC NREM sleep, as shown in Fig. 3l–m) and CFC recall-specific ensemble cells (Extended Data Fig. 4b) during post-CFC NREM sleep but not in the post-CFC awake state or during REM sleep (Extended Data Fig. 4c–e). These results imply that NREM sleep likely acts as an intermediate phase that continuously links NREM-dependent co-reactivation dynamics and prospective REM-dependent schema processing.

Furthermore, compared with the neutral group, the SoD group displayed increased co-reactivation between SI avoidance-related cells and CFC freezing-related cells (Fig. 3p) during post-CFC sleep but not during wakefulness (Fig. 3q–r), further suggesting the importance of sleep in strengthening neural representations underlying future biased emotional responses.

Together, these results suggest that in a negative emotional state, neuronal representations of emotional experiences are continuously processed and evolve flexibly during offline sleep after a new emotional experience.

### Causal Role of SoD-Emergent Ensembles and Sleep Dynamics in Negativity Bias

We next investigated whether emotional state-dependent negative ensembles emerging during the SoD session causally encode generalized emotional states. A mixture of AAV9-TRE3G-Cre and AAV9-CaMKII-DIO-hM4Di-mCherry was bilaterally injected into the dorsal hippocampus of cFos-tTA transgenic mice. This approach enabled us to tag ensembles that were active on the last day of the SoD (or neutral) sessions with inhibitory chemogenetic receptors for future anxiety experiments (Fig. 4a–c). After 48 h of incubation for full DREADD expression on the tagged ensembles, we first assessed the baseline emotional state of the mice via an open field test (OFT) and measured the time spent in the centre zone as an indicator of anxiety. Compared with mice injected with saline (SoD-Veh), mice in which SoD-tagged ensembles were silenced (SoD-J60) explored the centre more, indicating reduced anxiety (Fig. 4d–e). Inhibiting neutral-tagged ensembles (Neutral-J60) did not differ from the saline-injected Neutral-Veh group, confirming the specificity of the SoD ensembles. Additionally, an elevated plus-maze (EPM) test was performed 24 h after the OFT, revealing that compared with the SoD-Veh mice, mice in which the SoD ensembles were suppressed explored more in the open arms, which was not observed in neutral comparisons (Fig. 4f–g). These results suggest that SoD-emergent ensembles encode generalized emotional states primed by episodic experiences, thereby facilitating negativity bias.

**Fig. 4.**
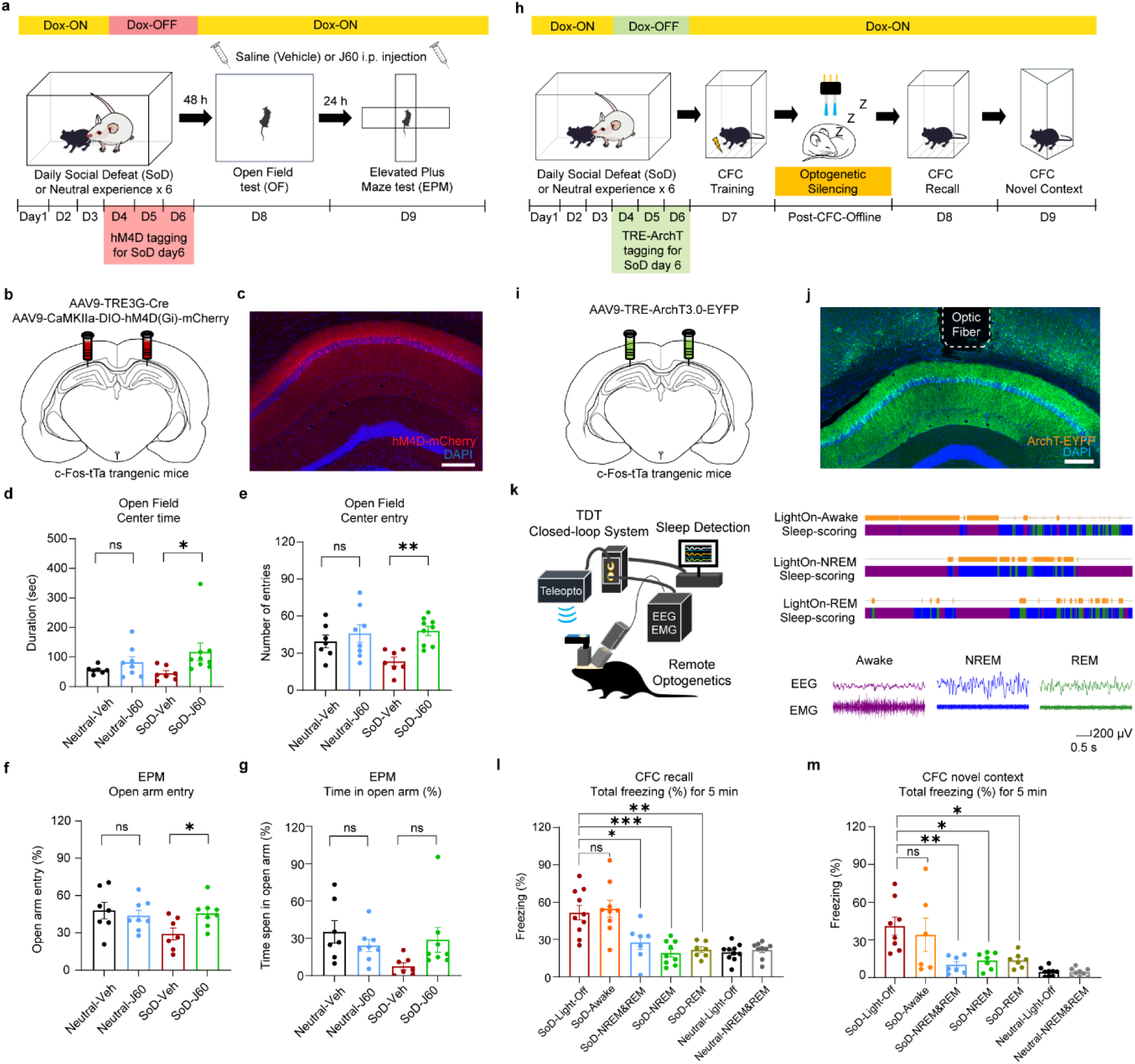
Chemogenetic and offline optogenetic silencing of SoD-tagged ensembles in the dorsal hippocampus decreases the negative emotional state and prevents emotional bias in future conscious experiences. **a.** Schematic of the timeline for chemogenetic silencing of SoD-tagged ensembles in the open field (OF) and elevated plus-maze (EPM) tests. **b–c.** Activity-dependent expression of inhibitory hM4D receptors on tagged SoD ensembles in the dorsal hippocampus. Scale bar, 250 µm. **d.** Total time spent in the centre of the OF in chemogenetic silencing experiments: each mice in the neutral and SoD groups was injected with either saline as the vehicle group (Veh) or the 3^rd^ generation of the DREADD agonist J60 (J60) as the drug group. (Neutral-Veh *n* = 7, Neutral-J60 *n* =8, SoD-Veh *n* = 7, SoD-J60 *n* = 9 mice; two-way ANOVA; interaction *p* = 0.2699; *F*(1, 27) = 1.269; Šídák’s multiple comparisons test). **e.** Total number of entries in the centre of the OF arena (two-way ANOVA, interaction *P =* 0.0878; *F*(1, 27) = 3.138; Šídák’s multiple comparisons test). **f.** Percentage of open arm entries in the EPM test (Two-way ANOVA, interaction *P* = 0.0452, *F*(1, 26) = 4.429, Šídák’s multiple comparisons test). **g.** Percentage of time spent in the open arms in the EPM test (Neutral-Veh *n* = 7, Neutral-J60 *n* =8, SoD-Veh *n* = 7, SoD-J60 *n* = 8 mice; two-way ANOVA; interaction *P* = 0.0374; *F*(1, 26) = 4.810; Šídák’s multiple comparisons test). **h.** Schematic of the timeline of optogenetic silencing of SoD-tagged ensembles during the post-CFC offline period and subsequent CFC recall tests and novel context exploration session. **i–j.** Activity-dependent expression of inhibitory archaerhodopsins (ArchT3.0) on the tagged SoD ensemble in the dorsal hippocampus and optic fibre location. Scale bar, 250 µm. **k.** Schematic of the closed-loop optogenetic silencing experiments during the first two hours of post-CFC offline period and example traces of EEG and EMG recordings during three offline sessions. **l.** Total freezing percentage (%) in the CFC recall session among the optogenetic experiment groups (SoD-Light-Off *n* = 10, SoD-Awake *n* = 9, SoD-NREM&REM *n* = 7, SoD-NREM *n* = 9, mice, SoD-REM *n* = 7, Neutral-Light-Off *n* = 10, Neutral-NREM&REM *n* = 9 mice, Brown–Forsythe ANOVA test, interaction *P* <0.001, *F*(4, 27.88) = 11.39, Dunnett’s T3 multiple comparisons test). **m.** Total freezing percentage (%) during the CFC novel context session among the optogenetic experiment groups (SoD-Light-Off *n* = 8, SoD-Awake *n* = 6, SoD-NREM&REM *n* = 7, SoD-NREM *n* = 7, mice, SoD-REM *n* = 7, Neutral-Light-Off *n* = 9, Neutral-NREM&REM *n* = 9 mice, Kruskal‒Wallis test, *P* = 0.0053, Dunn’s multiple comparisons test). Data are presented as the mean ± s.e.m. Statistical significance is expressed as follows: ns, not significant; * *P* < 0.05; ** *P* < 0.01; *** *P* < 0.001.

Next, we explored similar negativity representations in other higher-order areas, such as the neocortex, which is associated with schema formation^36^, affective processing^37–40^, inferential learning^18,20,41^, and fear^42,43^. We injected retrograde AAV2-Cre-EGFP into the dorsal hippocampus and AAV9-TRE-DIO-hM3-ALFA into the prefrontal cortex (PFC) of cFos-tTA mice, tagging SoD-specific PFC ensembles projecting to the dorsal hippocampus (Extended Data Fig. 5a–b). After a single acute SoD-ensembles were tagged, the activation of these PFC ensembles resulted in increased anxiety, as the SoD-J60 group showed decreased open arm exploration compared with the SoD-Veh and neutral groups (Extended Data Fig. 5c–d). Moreover, the activation of tagged PFC ensembles during the post-CFC training period induced greater freezing in the next-day recall session (Extended Data Fig. 6e–f). These data indicate that emotional state-dependent ensembles encoding negativity emerge in higher-order areas such as the PFC and generate persistent bias via postexperience activation.

We next explored whether SoD ensemble dynamics during sleep causally drive future negativity bias. AAV9-TRE-ArchT3.0 was bilaterally injected into the dorsal CA1 region of cFos-tTA mice to tag SoD ensembles with inhibitory opsins, followed by optic fibre implantation to enable wireless optogenetics (Teleopto) combined with EEG/EMG electrode recordings (Fig. 4h–j). This enabled the implementation of closed-loop optogenetic perturbations via the Tucker–Davis Technologies (TDT) system, which is used to score sleep states in real time on the basis of EEG/EMG recordings (Fig. 4k). For offline optogenetic inhibition, we tested the following: light-off control, inhibition while awake, inhibition during NREM sleep, inhibition during REM sleep, or inhibition during all sleep periods (NREM and REM) during the first two hours of the post-CFC offline periods. In the CFC recall session the following day, the SoD mice robustly displayed reduced freezing responses only in the sleep-silenced groups (NREM, REM, or both), whereas the awake-silenced group (SoD-Awake) presented freezing levels comparable to those of the SoD-light-off control group (Fig. 4l and Extended Data Fig. 6a).

To confirm the causal effect of sleep neural dynamics on biased emotional responses in general, we further compared freezing levels in a novel context one day after the recall session. Similarly, among the SoD mice, lower freezing was observed in the sleep-silencing groups but not the awake-silencing group (Fig. 4m). The lack of differences in the proportion of ArchT3.0-tagged cells between the neutral and SoD groups and the comparable light-on durations between the awake and NREM sessions (with the light-on duration in the REM session being the shortest) suggest that the effects of optogenetic inhibition likely stem from specific reactivation dynamics rather than the number of tagged cells or inhibition length (Extended Data Fig. 6b–c).

Together, these results demonstrate that SoD-emergent neuronal representations encode experience-dependent negativity and that dynamics in sleep, but not in the awake state, drive biased emotional responses in future conscious states. This highlights the biological importance of sleep-dependent processes in sustaining negativity bias.

### Sleep Neural Dynamics Shape Future Emotional Representations and Bias

Finally, we investigated how distinct neural representations embedded within conjunctive multimodal emotional experiences evolve and interact during sleep, thereby influencing emotional perceptions of past and future experiences. We implemented a trace fear conditioning (TFC) paradigm in which a long interval exists between the tone and aversive stimuli (Extended Data Fig. 7a). The TFC paradigm is known to require more conscious effort depending on higher-order functions in the neocortex and hippocampus than the classical delayed fear conditioning paradigm^44–46^, and this paradigm enabled us to identify the neural representations of precisely timed, cue-induced emotional states; furthermore, compared with the context-based CFC paradigm, this approach offers better temporal resolution. The CS that was previously paired with mild electric shocks (US) induced weak negative memories in the control recall (Recall 1) session on day 3 (Extended Data Fig. 7b–d). After retraining (Training 2) on day 4, followed by either repeated neutral or SoD experiences (day 5-10), we introduced a novel tone (ambiguous tone as a generalizing stimulus, GS) test session (day 11) one day prior to recall session 2 (Recall 2) on day 12 to avoid potential extinction effects. Compared with the neutral group, the learned CS induced greater freezing in the SoD group during both the tone and trace periods in recall session 2 on day 12 (Extended Data Fig. 7e–g). Novel tones, which were not paired with shock, also increased freezing in the SoD group on day 11 (Extended Data Fig. 7h–k). Thus, the SoD experience drives negatively biased interpretations of both past weak salient stimuli and novel stimuli in future conscious states.

We then conducted longitudinal Ca^2+^ imaging with EEG/EMG to track neural representation dynamics during online and offline periods (Fig. 5a) in SoD mice. We measured the similarity in the neural representations by quantifying the relative cosine angles between the population activity vectors (PVs) in vector space (Fig. 5b–e and Extended Data Fig. 8a–b). When the PV of recall session 2 with CS^+^ (D12 as day 12) was taken as a reference, the PVs of the post-SoD sleep session (D10, either when SoD cells reactivated or co-reactivated) exhibited significantly narrower cosine angles than did the PVs of recall session 2 with CS^−^ (D12) (Fig. 5b–c), the SoD day 6 session and post-SoD awake session (D10) (Fig. 5d–e), and the pre-SoD habituation session (D1) and training session 2 with CS^+^ (D4) (Extended Data Fig. 8a–b). For the global structure, we applied multidimensional scaling (MDS) to cross-session cosine distance matrix, projecting the PVs into 3D space. The PVs of the post-SoD sleep session clustered closest to the PV of recall session 2 with CS^+^ (Fig. 5f and Extended Data Fig. 8c). These results suggest that in a SoD-driven negative emotional state, sleep reshapes the representations of past salient experiences towards more negative direction, thereby leading to biased emotional interpretation in the future.

**Fig. 5.**
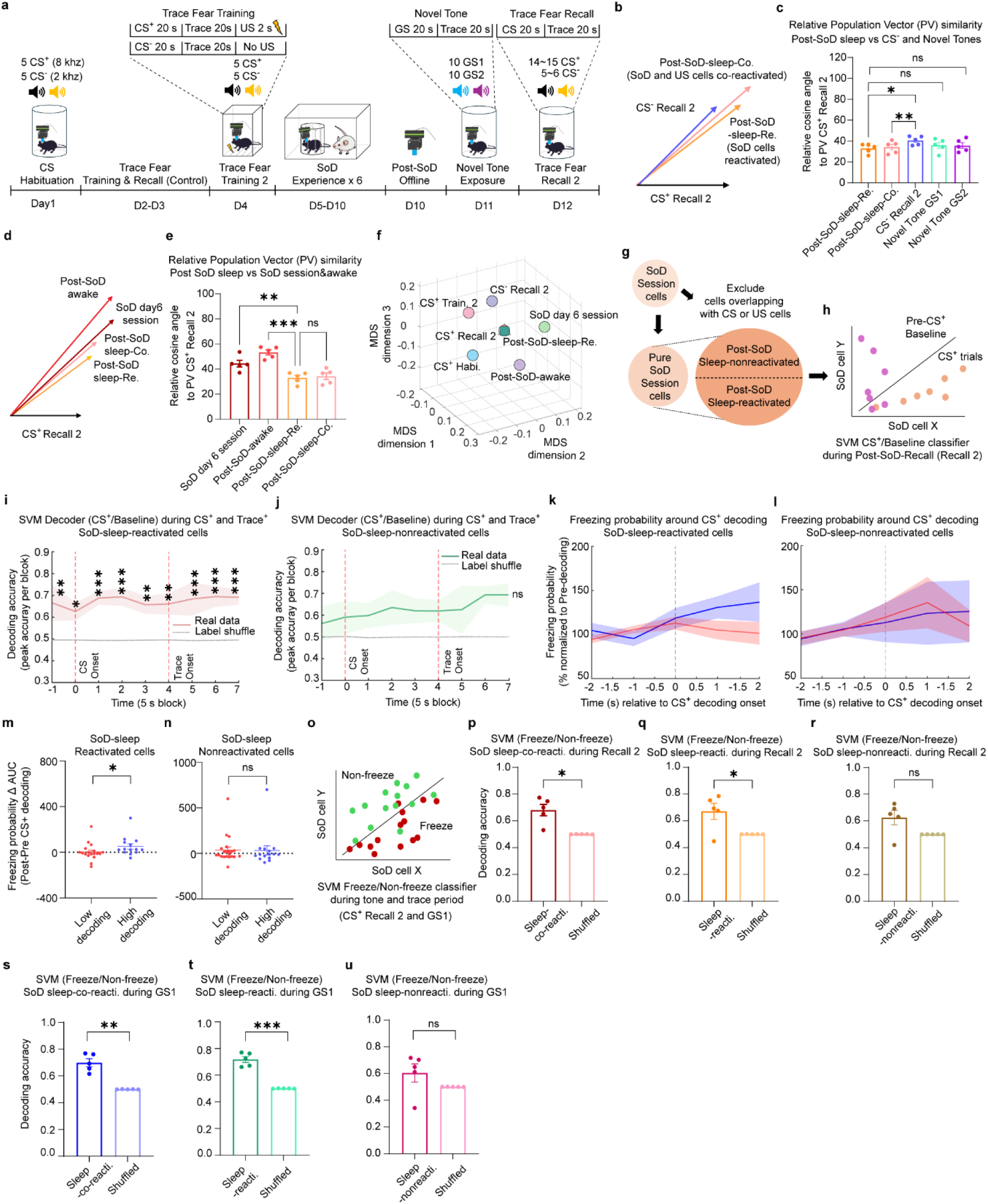
Negative-state sleep population dynamics guide evolving negative representations of past and future experiences. **a.** Schematic of the Ca^2+^ imaging recording schedule of the trace fear conditioning-SoD-Trace fear recall and novel tone exposure paradigms. **b.** 2D plot of the relationship of population activity vectors (PVs) between neural representations of CS^+^ trials in the recall period after SoD (CS^+^ Recall 2 on D12) as a reference and neural representations of CS^−^ trials in the recall period after SoD (CS^−^ Recall 2 on D12) and neural representations during sleep after the last day of SoD (on D10): when SoD neurons were reactivated (post-SoD-sleep-Re.) or when SoD neurons were co-reactivated with US neurons. **c.** Relative cosine angle between the reference PV (CS^+^ Recall 2) and the PVs of the post-SoD sleep period, CS^−^ recall session 2 and the novel tone session on D11 (*n* = 5 mice; one-way RM ANOVA with correction; *P* = 0.0103; *F* (1.459, 5.838) = 12.21; Dunnett’s multiple comparisons test). **d.** 2D plot of the relationship between the PVs of CS^+^ recall session 2 and the PVs of the SoD day 6 session, post-SoD awake session and post-SoD sleep session on D10. **e.** Relative cosine angle between the reference PV (CS^+^ Recall 2) and the PVs of the SoD session and post-SoD offline session (awake and sleep) (one-way RM ANOVA, *P* < 0.001; *F*(3, 12) = 21.29; Dunnett’s multiple comparisons test). **f.** Multidimensional scaling (MDS) of the cross-session cosine distance matrix from the PVs. Square-shaped (Post-SoD-sleep-Co.), triangular-shaped (post-SoD-sleep-Re.) and circular-shaped (CS^+^ Recall 2) PVs overlap in the middle. **g.** Schematic of pooling sleep-active SoD populations that do not overlap with CS- or US-related neurons of training 2 session before the SoD experience. **h.** Illustration of the support vector machine (SVM) decoder for classifying the CS^+^ and pre-CS^+^ baselines during the post-SoD recall period (Recall 2). **i.** Averaged peak decoding performance of the SoD-sleep-reactivated population per 5-second block of classifying CS^+^ from the pre-CS^+^ baseline in recall session 2 during the CS^+^ and trace^+^ trials compared with the shuffled datasets (*n* = 5 mice, 30 repeats of 10-fold cross validation per second of the trials, 1000 iterations of label shuffled surrogate datasets; two-way ANOVA, interaction *P* = 0.9891, *F* (8, 72) = 0.2053, Šídák’s multiple comparisons test), **j.** From the SoD-sleep-nonreactivated population with shuffled datasets (*n* = 5 mice; two-way ANOVA, 30 repeats of 10-fold cross validation per second of the trials; 1000 iterations of the label-shuffled surrogate datasets; interaction *P* = 0.9252; *F*(8, 72) = 0.3852; Šídák’s multiple comparisons test). **k.** Plot of the average freezing probability of all recorded mice around timepoints when the CS^+^ decoding accuracy is high (blue, top 20% decoding performance) and when the CS^+^ decoding accuracy is low (red, bottom 20% decoding performance) across all combined CS^+^ and trace^+^ trials from SoD-sleep-reactivated neurons, **l.** From SoD-sleep-nonreactivated neurons. **m.** Change in the area under the curve (delta AUC) of the freezing probability before CS^+^ decoding (pre-CS decoding) and after CS^+^ decoding (post-CS decoding) of SoD-sleep-reactivated neurons. Low decoding indicates timepoints where CS^+^ decoding accuracy was low, whereas high decoding indicates timepoints where CS^+^ decoding accuracy was high across the trials (*n* = 5 mice; low decoding accuracy, *n* = 18 timepoints; high decoding accuracy, *n* = 14 timepoints; Mann–Whitney *U* test, *P* = 0.0215). **n.** Change in the AUC of the freezing probability from pre- to post-CS decoding onsets of SoD-sleep-nonreactivated neurons (*n* = 5 mice; low decoding accuracy, *n* = 21 timepoints; high decoding accuracy, *n* = 15 timepoints; Mann–Whitney *U* test, *P* = 0.7453). **o.** Schematic of the SVM decoder for classifying freezing and nonfreezing behavioural states (SVM Freeze/Non-Freeze Decoder) during the CS^+^ and Trace^+^ periods of recall session 2. **p.** Freeze state decoding performance of SoD-sleep-co-reactivated neurons (Sleep-co-reacti.) (*n* = 5 mice; unpaired t test with correction; *t* = 4.216; *P* = 0.0135), **q.** Same as **p** for SoD-sleep-reactivated neurons (Sleep-reacti.) (Unpaired t test with correction, *t* = 2.875, *P* = 0.0452) and **r.** SoD-sleep-nonreactivated neurons (Sleep-nonreacti.) (Unpaired t test with correction, *t* = 2.324, *P* = 0.0807) with shuffled datasets. **s.** Freeze state decoding performance of SoD-sleep-co-reactivated neurons (Sleep-co-reacti.) during novel tone 1 (GS1-tone and trace periods combined) trials (*n* = 5 mice; unpaired t test with correction, *t* = 6.671; *P* = 0.0026), **t.** Same as **s** for SoD-sleep-reactivated neurons (Sleep-reacti.) (Unpaired t test with correction, *t* = 9.663, *P* < 0.001) and **u.** SoD-sleep-nonreactivated neurons (Sleep-nonreacti.) (Unpaired t test with correction, *t* = 1.524, *P* = 0.2021) with shuffled datasets. Data are presented as the mean ± s.e.m. Statistical significance is expressed as follows: ns, not significant; * *P* < 0.05; ** *P* < 0.01; *** *P* < 0.001.

Then, we examined whether SoD cells that were active during sleep functionally compute negative stimuli in future experiences. We first pooled SoD session-related cells, excluding any cells that overlapped with prior CS-related or US-related cells, and then extracted subsets that were reactivated (sleep-reactivated) or not (sleep-nonreactivated) during post-SoD sleep (Fig. 5g). We trained linear support vector machine (SVM) decoders with either of these two types of cells to classify early CS^+^ trials from pre-CS^+^ baselines in the recall 2 session (D12) (Fig. 5h). Sleep-reactivated SoD cells could successfully decode CS^+^ trials, including tone and trace periods (Fig. 5i), whereas non-reactivated SoD-cells could decode CS^+^ trials only during the trace period, albeit not significantly (Fig. 5j), indicating the differential roles of populations that were active or not active during sleep. Within the populations that were active during sleep, we observed comparable proportions of NREM- and REM-selective cells, suggesting the functional importance of both sleep processes (Extended Data Fig. 9a).

To link CS decoding to biased negative behavioural responses, we quantified the freezing probability around timepoints with high (top 20% decoding performance) and low (bottom 20% decoding performance) decoding accuracy during these trials (Fig. 5k–l and Extended Data Fig. 9b-0d). The quantification of the changes in freezing probability from pre- to post-CS^+^ decoding onset revealed that compared with low-decoding-accuracy timepoints, freezing increased significantly post-onset for high-decoding-accuracy timepoints of sleep-active SoD cells (Fig. 5k, m), whereas no difference was observed for non-reactivated cells (Fig. 5l, n). These findings suggest that sleep-engaged SoD subsets encode aversive stimuli that are predictive of biased emotional responses.

To further test whether sleep-active populations directly decode biased negative emotional behaviour, we trained an SVM to classify freezing and non-freezing behaviours across CS^+^ and trace trials (Fig. 5o and Extended Data Fig. 10a–c). Sleep-active SoD cells, either reactivated (Sleep-reacti.) or co-reactivated with the previous US (Sleep-co-reacti.) cells, decoded freezing behaviour successfully (Fig. 5p–q), whereas non-reactivated cells (Sleep-nonreacti.) showed lower performance comparable to shuffle levels (Fig. 5r). Finally, we assessed the ability of the cells to decode freezing behaviour during trials with novel GS tones. Sleep-active, but not non-active, SoD cells decoded freezing behaviours during trials with novel ambiguous stimuli (Fig. 5s–u). Thus, sleep-active SoD cells compute generalizable negativity beyond memory-related experience.

Together, these results demonstrate that emotional state-dependent neural dynamics during sleep shape the emotional representations conveyed to future conscious experiences by recruiting subpopulations of ensembles associated with stressful negative experiences to functionally compute biased perception and stimulus-guided emotional responses. This highlights the role of offline sleep periods in perpetuating and reshaping negativity representations.

## Discussion

Our findings reveal that stress-driven negative emotional states lead to the robust emergence of semantic-like neuronal representations of negative emotions in hippocampal cellular subnetworks that biologically correlate with biased emotional responses. Our results further demonstrate how integrative neural computations of past and new emotional experiences during sleep reshape negative schema-like representations. Neural spaces during sleep ensure the persistence of negativity, with emotional schemas during subconscious sleep states internally updated and later translated to conscious future experiences.

Our longitudinal Ca^2+^ imaging data revealed neural signatures of semantic-like negativity representations that emerged during repeated negative experiences at both the population and single-cell levels. Mice in the negative states exhibited specific neural ensembles that responded to various negative emotional experiences across distinct timepoints and environments, suggesting that these populations compute conceptual aspects of negative emotion by abstracting its gist from episodic experiences. Mechanistically, our results suggest that representations of negativity bias evolve by enhancing functional interactions among subtypes of negative emotion-related cells, which enables the flexible formation of neural structures serving as cognitive maps of emotional states. The lack of significant differences between the negative state and neutral groups in terms of overlapping cell proportions among negative emotion subtypes indicates that biased responses likely arise from the dynamic coordination of independent subsets of cells rather than mechanisms relying on overlapping populations, which are typically associated with memory linking^47–50^. Remarkably, sleep-active SoD cells, which are distinct from CS and US representations, can independently decode aversive CS^+^ and freezing behaviours. This provides novel mechanistic insight into negativity bias, with these cell populations harbouring information that affects future negative behaviour.

The dorsal hippocampus in rodents (posterior in humans) is involved in complex multimodal episodic processing, whereas ventral (anterior) regions regulate emotion-related behaviours^51,52^. Our Ca^2+^ imaging and chemogenetic results show that the dorsal hippocampus generates experience-dependent emotional representations reflecting semantic-like negativity. This finding is consistent with findings that artificial manipulation of defeat-tagged dorsal hippocampal engrams modulates social avoidance^53^. These results thus imply that episodic ensembles store information related to the emotional state that is embedded within conscious experiences. Unlike the ventral hippocampus and amygdala, which contain innate-hardwired cells associated with anxiogenic environments^54,55^, fear^56,57^ or valence^58^, our findings suggest that the dorsal hippocampus may lack such defaults but instead likely stores higher-order emotional concepts via conjunctive episodic units^59^, thereby reflecting the ability of humans to consciously recall emotions tied to autobiographical memories^60^. Furthermore, our data suggest that the dorsal hippocampus might engage in more gradual/perpetual emotional processing in an emotional state-dependent manner, which aligns with the finding that trait anxiety is associated with the posterior hippocampus in humans.^61^

Higher-order cognitive functions involve complex interregional coordination^62^; thus, similar negativity representations likely exist elsewhere, as our data suggest that experience-dependent prefrontal populations modulate the corresponding bias. This suggests the future direction of the functional specification of cognitive emotional processing areas and their coordination with known subcortical limbic systems across online-to-offline phases. We speculate that neural coding of long-lasting subjective emotions—likely combining subconscious autonomic and conscious aspects^3,5,63^—requires bidirectional bottom-up to top-down processing, with gaps in synaptic and circuit mechanisms warranting exploration. Previous rodent studies have reported off-task reactivation of a single aversive experience during awake^64^ or sharp-wave ripple states^11,65^. Here, we discovered that sleep orchestrates interactions among multiple emotionally congruent experiences and reshapes negative schema-like representations, which perpetuate to future experiences. The optogenetic inhibition results confirm that awake processes are insufficient for ensuring causal future bias, highlighting the unique functional role of offline sleep periods compared with offline awake processing periods. We speculate that resting-state neural activity in the awake state, driven by conscious attention, processes highly salient emotional stimuli and new experiences via focused reactivation to promote rumination, whereas during sleep, which does not involve attentional constraints, facilitates broad semantic evaluation of subjective emotions for the future through spontaneous, integrative reactivations. In this sleep-driven process, the cognitive definition of subjective negativity may be dynamically adjusted, broadening or constricting the boundaries of how autobiographical experiences are emotionally categorized and interpreted. We speculate that this homeostatic-like adjustment may be disrupted in individuals with mood disorders, leading to persistent negative overgeneralization through excessive broadening of these boundaries.

Furthermore, we observed distinct neural dynamics of stressed mice during sleep: co-reactivation of distinct emotional experiences in NREM sleep and activation of future-conveying negative schema in REM sleep. NREM sleep drives co-reactivation, likely resulting in the extraction of abstract representations of negativity from distinct emotional experiences. In contrast, in REM sleep, which is linked to integration and generalization^66^, the updated neural template of the proto-negative schema-like representation may be stabilized. Both phases are causally important for future bias on the basis of our optogenetic data (Fig. 4l–m). Future studies should explore the differential and interdependent roles played by NREM and REM sleep in the creation and interpretation of emotional representations.

Population vector and MDS analyses revealed similar neural representations between post-SoD sleep sessions and future aversive CS^+^ sessions, highlighting the predictive role of sleep in preparing for future negative interpretations without conscious effort. Sleep-active SoD populations, which can decode aversive CS and freeze responses across both learned and novel tones, demonstrate continuous reshaping of generalized negativity representations, extending beyond fear memory-specific components through multi-experience integration. Sleep may shape predictive emotional representations from integrating congruent experiences, thereby guiding future biased interpretations.

Taken together, our findings provide novel biological evidence for emotional schema-like representations that reflect semantic negativity in cognitive memory networks, offering primitive support for theoretical models of higher-order cognitive encoding in emotional interpretation. We offer a new framework in which subconscious sleep dynamics may serve as a cognitive platform for evaluating subjective emotions, guiding emotional inception, i.e., predefining perspectives that are later used consciously. Our findings provide mechanistic insights into long-term bias in intense negative states, thereby suggesting a new perspective on cognitive emotional processing and novel therapeutic scopes for mood disorders, along with current neuropharmacological treatments and cognitive behavioural therapies.

## Methods

### Animals

All animal procedures complied with the guidelines of the National Institutes of Health (NIH) and were approved by the Animal Care and Use Committee of the University of Toyama. Naïve wild-type C57BL/6J male mice (SLC Japan Inc.) and c-Fos-tTA transgenic mice (Mutant Mouse Regional Resource Center, stock number: 031756-MU) were maintained on a 12-hour light-dark cycle with food and water provided ad libitum. For c-Fos-tTA transgenic mice, food containing 40 mg/kg doxycycline (Dox) was provided until the activity-dependent tagging experiments and switched to 1000 mg/kg Dox after cell tagging. All mice, aged 16–32 weeks, were used for behavioural and recording experiments.

### Viral constructs

For longitudinal *in vivo* Ca²⁺ imaging experiments, we used a recombinant adeno-associated virus (AAV) vector, AAV9-hSyn-jGCaMP7b (Janelia variant, Addgene: #104489, viral titer: 9.77 × 10¹³ vg/mL). For inhibitory chemogenetic DREADDs experiments, we subcloned the hM4D(Gi)-mCherry fragment from pAAV-CaMKIIa-hM4D(Gi)-mCherry (Addgene: #50477) to replace the hM3D(Gq)-mCherry fragment in our custom-produced pAAV-CaMKIIa-DIO-hM3D(Gq)-mCherry (originally from pAAV-hSyn-DIO-hM3D(Gq)-mCherry, Addgene: #44361) to create pAAV-CaMKIIa-DIO-hM4D(Gi)-mCherry (viral titer: 9.58 × 10¹² vg/mL). For activity-dependent tagging of inhibitory DREADDs receptors, we used pAAV-TRE3G-Cre-WPRE (viral titer: 1.28 × 10¹⁴ vg/mL), as previously described^68^. For activity-dependent tagging of excitatory chemogenetic DREADDs circuit experiments, we subcloned hM3D(Gq)-ALFA fragment (hM3D(Gq) from Addgene: #50460 and ALFA sequence from the previous report^69^) from pEX-A2J2-DIO-hM3D(Gq)-ALFA to replace ChR2(T159C)-ALFA fragment from our custom-produced pAAV-TRE3G-DIO-ChR2(T159C)-ALFA (ChR2(T159C) originally donated by Dr. Karl Deisseroth) to finally create pAAV-TRE3G-DIO-hM3D(Gq)-ALFA (Eurofins Genomics, viral titer: 5.71 × 10¹⁴ vg/mL). For retrograde delivery of Cre from dorsal hippocampus, we purchased rAAV2-CAG-Cre-EGFP (SignaGen: #SL11604, viral titer: 1.10 × 10¹³ vg/mL). For optogenetic silencing experiments, we used pAAV-TRE3G-ArchT3.0-EYFP (viral titer: 4.4 × 10¹³ vg/mL), as previously described^70^. Except for the purchased rAAV2 virus, all recombinant AAV vectors were packaged as serotype AAV9. All virus stocks were stored at −80°C until stereotactic surgeries.

### Stereotactic virus injection

Mice were anesthetized with an intraperitoneal injection of a mixture containing 0.75 mg/kg medetomidine (Domitor; Nippon Zenyaku Kogyo Co, Ltd., Japan), 4.0 mg/kg midazolam (Fuji Pharma Co., Ltd., Japan), and 5.0 mg/kg butorphanol (Vetorphale; Meiji Seika Pharma Co., Ltd., Japan). During surgery, mice were placed on a stereotactic apparatus with a heating pad to maintain body temperature. After marking the target coordinates and drilling a hole at the coordinate area, the AAV virus in a mineral oil-filled glass needle attached to a 10 µL Hamilton syringe (80030, Hamilton, USA) was injected using an automated motorized microinjector IMS-20 (Narishige, Japan). Post-surgery, 1.5 mg/kg atipamezole (Antisedan; Nippon Zenyaku Kogyo Co., Ltd., Japan), an anti-sedative, was administered via intramuscular injection. Saline and Ringer’s solution (0.5 mL/mouse, i.p.; Otsuka., Japan) were injected to aid recovery. All procedures were conducted as previously described^71,72^. Briefly, for chemogenetic silencing of the dorsal hippocampus, 500 nL of a virus mixture of AAV9-TRE3G-Cre and AAV9-CaMKIIa-DIO-hM4D(Gi)-mCherry (1:2 ratio) was bilaterally injected into the dorsal hippocampal CA1 (AP: −2.2, ML: ±1.2, DV: −1.4 from Bregma). For optogenetic silencing, 500 nL of AAV9-TRE3G-ArchT3.0-EYFP was bilaterally injected into the hippocampal CA1 area. For chemogenetic activation in the prefrontal cortex (PFC), 500 nL of rAAV2-CAG-Cre-EGFP was bilaterally injected into the same hippocampal coordinates, while 500 nL of AAV9-TRE3G-DIO-hM3D(Gq)-ALFA was bilaterally injected into the ACC (and borderline region with M2) area (AP: +2.2, ML: ±1.2, DV: −1.3 from Bregma) in the same mouse. For *in vivo* Ca²⁺ imaging, 500 nL of AAV9-hSyn-jGCaMP7b was unilaterally injected into the dorsal hippocampal CA1. All virus injections were followed by maintaining the glass injection tip in place for an additional 5 minutes before gently withdrawing it from the brain tissue. Viruses were injected at a rate of 100 nL/min.1.

### Optogenetic fiber and EEG/EMG implantation

After injecting AAV9-TRE3G-ArchT3.0-EYFP, a dual-LED cannula (fiber diameter: 500 µm, fiber length: 3.1 mm, bilateral, 590 nm) of a wireless optogenetic system^72,73^ (Teleopto; Bio Research Center, Japan) was implanted above the injected CA1 area (AP: −2.2, ML: ±1.2, DV: −1.1 from Bregma) on the same day. Concurrently, EEG/EMG electrodes were implanted as previously described^20,71^. Briefly, EEG electrode-attached screws were placed in the skull over the parietal cortex, posterior to the optic fiber locations while ground and reference electrode-attached screws were placed over the right and left cerebellar cortex, respectively. Two EMG wires were implanted into the neck muscle just below the skull. Dental cement (Shofu Inc, Kyoto, Japan) was used to secure the implanted optic fibers and EEG/EMG electrodes.

### Inscopix GRIN lens and EEG/EMG implantation

Following >1 week after AAV9-hSyn-jGCaMP7b injection, mice were anesthetized and fixed on a stereotactic apparatus. Following craniotomy above the injected area, the neocortex and corpus callosum above the alveus were aspirated using a 26-gauge blunted needle tip with continuous saline irrigation. After bleeding ceased, the GRIN lens (Inscopix, CA, diameter: 1.0 mm, length: 4 mm) was slowly implanted using custom-made forceps attached to a manipulator (Narishige, Japan) until it contacted the tissue surface. Emulsified bone wax was applied to fill the gap between the craniotomy and the lens, followed by dental cement to secure the lens. Subsequently, similar to the optogenetic procedure, an EEG recording electrode screw was implanted into the skull over the parietal cortex, with reference and ground electrodes placed above the cerebellar cortex, and EMG electrodes implanted in the neck muscle. Additional dental cement was applied to secure the EEG/EMG setup and GRIN lens. A black-colored surface EP tube was placed over the lens for protection until baseplate attachment. Two weeks later, baseplate installation was performed on the stereotactic apparatus after anesthesia. The baseplate was temporarily attached to a miniature microscope (nVista3, Inscopix, CA) with a loose screw and lowered above the implanted GRIN lens until a clear view of vasculature was observed via real-time recording on a PC monitor. After fine-tuning the focus, black-colored dental cement mixed with 5% carbon powder (#64164, Sigma) was applied to fix the baseplate above the lens center. Once the cement had hardened, the baseplate was detached from the miniature microscope, and mice were returned to their homecage for at least two weeks before *in vivo* Ca²⁺ imaging experiments.

### Behavioural experiments

In prior to all behavioural experiments, mice were gently handled for three days and were habituated to the resting space for 15 minutes before transferring to the experimental room. Mice were also habituated inside the soundproof room for 30 minutes. For all chemogenetics and optogenetics experiments, mice with only unilateral, off-target to no viral expressions or damaged brain tissues in the target area were excluded from the data.

### Repeated social defeat stress (SoD) protocol

We implemented a standard social defeat stress (SoD) protocol^31,74^ with modifications, as previously described. After screening aggressive retired breeder male ICR mice (Crl:CD1 (ICR), Charles River Inc.), experimenter C57BL/6J mice (wild-type naïve or c-Fos-tTA transgenic) were placed in the homecage of aggressive ICR mice. This resident-intruder protocol allowed resident ICR mice to attack intruder B6 mice. After 2–3 minutes, including at least 10 bouts of aggressive attack, defeated experimenter B6 mice were returned to their homecage until the next day. The SoD protocol was conducted for six consecutive days, with each experimenter B6 mouse encountering a novel aggressive ICR mouse daily. For Ca²⁺ imaging experiments, to prevent ICR mice from damaging the Inscopix microscope cable, we introduced an additional session after the direct attack, placing defeated B6 mice inside a wire-mesh cup in the homecage of aggressive ICR mice. This allowed minimal sensory contact with ICR mice while protecting the microscope cable during recording. Neural data on SoD day 1 and SoD day 6 were recorded while defeated B6 mice were in the wire-mesh cup. For the neutral experience group (Neutral), experimenter B6 mice freely explored a novel homecage without social stimuli for 3 minutes and were returned to their homecage. For Ca²⁺ imaging experiments of the neutral group, B6 mice were placed in a wire-mesh cup in a novel homecage without social stimuli after 3 minutes of free exploration. Separate cohorts were used for the SoD-social interaction (SI) and the SoD-Elevated plus maze (EPM) paradigms in Figure 1a-f. All cohorts in the SoD-SI paradigm went through subsequent contextual fear conditioning (CFC) of Figure 1g-k (the last cohort, consisting of 4 animals each in the neutral and SoD groups, went through the SI procedure without video recording due to a technical issue with the camera on the day of social interaction).

### Female social interaction (FSI) protocol

Each day, naïve wild-type male mice were placed in the homecage of a novel virgin wild-type female mouse (aged 13–20 weeks) for 2 hours and then returned to their homecage. This duration facilitated active social interaction without mating. Similar to the SoD protocol, mice encountered a new female mouse daily. As a control, male mice explored a novel homecage without social stimuli for 2 hours and were returned to their homecage.

### Social interaction (SI) test

The social interaction (SI) test was conducted as previously described^31,74^ with minor modifications. Briefly, experimenter B6 mice (after neutral or SoD experience) were placed in an open field arena (40 cm × 40 cm) and allowed to freely explore with an empty wire-mesh cage positioned at the center of one edge for 2.5 minutes. After the habituation session, a novel ICR mouse, not previously encountered during SoD, was placed in the wire-mesh cage, and experimenter B6 mice were allowed to freely explore and interact with the caged ICR mouse for 2.5 minutes. Social interaction time and time spent in opposite corners were quantified as measures of social avoidance, with the interaction zone defined as a rectangular area (20 cm × 10 cm) surrounding the wire-mesh cage, and opposite corners defined as square areas (10 cm × 10 cm) at each corner on the opposite side of the interaction zone. For Ca²⁺ imaging experiments, we conducted two consecutive 2.5-minute sessions with a novel conspecific B6 stranger mouse in the wire-mesh cage (SI-B6) and three consecutive 2.5-minute sessions with a novel ICR mouse in the wire-mesh cage (SI-ICR), with an inter-session interval of 2 minutes and 30 seconds. All behavioural data were analyzed using automatic video tracking systems (ÒHara & Co., Tokyo, Japan) and Ethovision XT 15 (Noldus).

### Contextual fear conditioning test (CFC)

For contextual fear conditioning (CFC), two context chambers were used: a square-shaped Context A and a triangular-shaped Context B, each with distinct wall patterns. One wall of each context was made of transparent Plexiglass, placed above a grid floor consisting of stainless-steel rods connected to an electric shock generator (Muromachi Kikai, Tokyo). For the training period, both neutral and SoD group mice were placed in Context A and allowed to explore for 2 minutes. After this habituation period, mice received three 0.4 mA electric footshocks (inter-stimulus interval: 60 seconds, 1.5 s per shock), followed by a 30-second incubation period before returning to their homecages. After 24 hours, mice were returned to Context A for a fear memory recall test. Freezing behaviour was measured for 5 minutes using automatic scoring software (Muromachi Kikai, Tokyo) connected to a video camera above the chamber. For novel context experiments, mice were placed in Context B with a different wall and grid floor pattern one day after the memory recall test, and freezing behaviour was measured. For positive experience-CFC experiments, a training protocol of three 0.25 mA footshocks (inter-stimulus interval: 60 seconds, 1.5 s per shock) was used. For Ca²⁺ imaging, optogenetic silencing, and chemogenetic activation experiments, a training protocol of two 0.25 mA footshocks (inter-stimulus interval: 60 seconds, 2 s per shock) was used to account for post-surgical stress and intraperitoneal injection effects in chemogenetic experiments. For optogenetic experiments, later cohorts of SoD (or Neutral)-CFC recall tests were also used for subsequent novel context tests a day after recall.

### Open field test (OF)

After 30 minutes of habituation in the experimental room, mice were placed in the center of an open field arena and allowed to freely explore for 10 minutes under 50 lux lighting. To measure anxiety-like behaviour, we quantified the time spent in the center zone, defined as a 2 × 2 square area located at the center of a 4 × 4 grid of subdivided squares covering the arena from EthoVision XT 15 analysis. For chemogenetic experiments, both neutral and SoD groups were intraperitoneally injected with either saline (control vehicle group) or J60 (JHU37160 dihydrochloride, Hello Bio, 0.1 mg/kg), a third-generation DREADD agonist (drug group), 10 minutes before the start of the open field test.

### Elevated plus maze test (EPM)

The elevated plus maze was established as a maze consisting of two types of arms: open arms (length: 25 cm, width: 5 cm) and closed arms (length: 25 cm, width: 5 cm, height of the walls: 16 cm). The maze was elevated 50 cm above the ground. Mice were placed at the intersection of the arms and allowed to freely explore for 10 minutes under 100 lux lighting. To measure anxiety-like behaviour, we used automatic video tracking systems (Ethovision XT 15 and ImageEP program^75^ based on the public domain NIH ImageJ program with the plugins^76^) to calculate the percentage of time spent in the open arms relative to the closed arms and the number of entries into the open arms. For chemogenetic experiments, the same procedures as the open field test were followed: mice were intraperitoneally injected with either saline or J60 solution 10 minutes before the start of the experiment.

### Trace fear conditioning test (TFC)

For trace fear conditioning, two contexts with distinct shapes, walls, and floor patterns were used: a circular-shaped chamber with a flat floor for tone habituation, memory recall, and generalization sessions, and a square-shaped chamber with a grid floor for conditioning (Muromachi Kikai, Japan). On the habituation day, mice were placed in the circular chamber and allowed 3 minutes of habituation. Then, mice were presented with either an 8 kHz or 2 kHz tone for 20 seconds, followed by a 60-second inter-trial interval, with each tone presented five times. On the training day (training 1 session as initial training and training 2 session for retraining), after 3 minutes of habituation in the square chamber, an 8 kHz tone served as the conditioned stimulus (CS^+^), followed by a 20-second trace period and a delivery of 2-second 0.25 mA electric footshock as the unconditioned stimulus (US). A 2 kHz tone served as the control stimulus (CS^−^), not paired with a US, with a 20-second trace period after tone termination. Each CS^+^ and CS^−^ trial was repeated five times with a 40-second inter-trial interval. For trace fear memory recall tests (recall 1 session before SoD and recall 2 session after SoD), mice were returned to the circular chamber and presented with a mixed order of 14 (14-15 for Ca^2+^ imaging) CS^+^ and 6 (5-6 for Ca^2+^ imaging) CS^−^ trials (inter-trial interval: 40 seconds after the 20-second trace period) following a 3-minute habituation period. For novel tone experiments post-SoD, 10 trials of each 6 kHz (generalizing stimulus, GS1) and 3 kHz (generalizing stimulus, GS2) tone were presented with a 40-second inter-trial interval and a 20-second trace period in the circular chamber. To prevent the fear extinction effect caused by the memory recall test before SoD experience, mice were retrained (Training 2) with the same training procedure the day after Recall 1. Moreover, the day after the SoD experience, we introduced a novel tone experiment on the first day before introducing Recall 2 to purely capture the intact negativity bias affected by SoD without weakening the original memory trace and stimulus-driven emotional response as much as possible. As a control session, a memory recall test session before Neutral/SoD experience (Recall 1) was performed to make sure that the neutral group and the SoD group have comparable learning, memory, and associated emotional response before the SoD experience.

### Sleep scoring and detection system

All EEG/EMG data during Ca²⁺ imaging and optogenetic experiments were recorded and monitored using the OpenEx Software Suite v2.32 (RX8-2, Tucker-Davis Technologies, USA), as previously described^20,77^ with minor modifications. EEG and EMG signals were amplified, filtered at 1–40 Hz and 65–150 Hz, respectively, and digitized at a 508.6 Hz sampling rate. Sleep stages were determined by real-time analysis of the EMG root mean square (RMS) value, EEG delta power (1–4 Hz) RMS, and EEG theta power (6–9 Hz) RMS. The EMG-RMS threshold was optimized for each mouse during a habituation period before experiments. Sleep and awake states were differentiated by the EMG-RMS threshold for three consecutive 4-second bouts. NREM sleep was defined as a delta/theta (D/T) ratio exceeding 1 for a consecutive 12-second period, whereas REM sleep was defined as a D/T ratio below 1. Real-time visual inspection of mouse movement and delta-dominant (0.5–4 Hz) or theta-dominant (4–9 Hz) waveforms on the monitor was performed alongside the automatic scoring system. Recorded EEG/EMG data were later exported and analyzed using custom MATLAB code. For spectrogram analysis, continuous EEG and EMG signals were notch-filtered at 55–65 Hz to remove line noise using a zero-phase IIR filter. Spectrograms were computed using multitaper spectral estimation as implemented in the Chronux toolbox function ‘MTSpectrogram’. Spectral power was normalized to the peak value within each recording channel.

### Closed-loop optogenetic silencing experiment

Prior to optogenetic silencing experiments, mice were habituated to the attachment of the wireless Teleopto optogenetic system and EEG/EMG cables for three days. During offline optogenetic perturbation experiments, the Teleopto system^73^, with a fully charged battery, delivered LED light controlled by TTL pulses from the EEG/EMG monitoring Tucker Davies Technology (TDT) system. Optical LED illumination (590 nm, continuous light, ∼3 mW output from the optic fiber tip) was automatically triggered by real-time EEG/EMG sleep scoring in a closed-loop manner. Mice were randomly assigned to experimental groups (light-off control, awake inhibition, or sleep inhibition), and optogenetic silencing was performed during the first 2 hours of post-CFC training after returning to the homecage. All groups were subsequently sacrificed and perfused for histological validation of ArchT opsin expression and optic fiber placement.

### *In vivo* Ca²⁺ imaging data processing and cell detection

All Ca²⁺ imaging neural data were processed using Inscopix Data Processing Software (IDPS, Inscopix, CA), as previously described^20,71,72^. Briefly, raw data from each session were concatenated into a single movie, spatially downsampled by a factor of two, and motion-corrected against a pre-selected reference frame with stable motion and clear fluorescence and blood vessel visualization. Video files with excessive motion or out-of-focus imaging were excluded from further analysis. Motion-corrected videos were transferred to Inscopix Mosaic software and temporally trimmed to isolate neural data from sessions of interest. Each day’s data was processed to calculate the change in Ca²⁺ fluorescence (ΔF/F), and the resulting data were concatenated into a single ΔF/F movie across all sessions. Cells were identified using our automatic sorting algorithm, HOTARU, as previously described^20,71,72^. Detected Ca²⁺ signals were extracted into a time × neuron matrix format. For neural data analysis, extracted Ca²⁺ data were 1-second binned and z-score normalized.

### PCA/ICA analysis

To extract neural cell ensembles, defined as co-firing cell populations active during behavioural sessions of interest, we applied a dimensionality reduction method combining Principal Component Analysis (PCA) and Independent Component Analysis (ICA), as previously described^20,78,79^ with minor modifications for tracking reactivation of multiple neural ensembles across longitudinal recording sessions. For extracting CFC-recall-specific ensembles, PCA was applied to z-score normalized neural data from the CFC recall session (CFC Recall). Principal components were pooled based on the Marchenko-Pastur bound, and ICA (fastICA) was subsequently applied to these components to extract independent neural sub-ensembles (cell assemblies). For each ensemble, neuronal weights were thresholded at 2 standard deviations (2 SD), and a projection matrix was formed with the diagonal set to zero to mitigate the influence of high-firing neurons on ensemble activity. These projection matrices were backprojected to other sessions to detect ensemble activity across days. Neural ensemble activation was defined as:

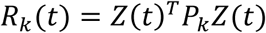

which *R*_*k*_(*t*) is the activation strength of the ensemble *k* at time *t*, *P* is the projector and *Z* is the other session(s) for backprojection.

To quantify mean ensemble activity per session, we calculated the activity rate, defined as the total number of ensemble activations above 2 SD divided by the duration of each corresponding session. CFC recall-specific ensembles were identified as those showing significantly higher activity rate (Wilcoxon rank-sum test) in the CFC recall session compared to the CFC pre-shock session (Pre-shock, a 2-minute habituation period before electric footshock). Ensembles without significant activity differences between CFC recall and CFC pre-shock sessions were classified as CFC recall-nonspecific ensembles. To compare the specificity of CFC recall-specific ensembles between neutral and SoD groups, the activity index of each CFC recall-specific ensemble per session was normalized by the mean activity rate of CFC recall-nonspecific ensembles per each corresponding session. Normalized CFC recall-specific ensemble activity index was compared between neutral and SoD groups. Pearson correlation was used for correlating the mean normalized activity rate of CFC recall-specific ensembles during the SI-ICR session to the total freezing level during the CFC recall session per mouse. For session-to-session comparisons within groups, the activity rate of CFC recall-specific ensembles was compared within each group (Neutral and SoD group), from pre-experience homecage to experience (sensory contact session after neutral or SoD episode) on day 6, from SI-B6 session to SI-ICR session, and from CFC pre-shock session to CFC post-shock session (CFC Post-shock, remaining session after the delivery of the first electric footshock) on CFC training day.

To assess correlations between averaged CFC recall-specific ensemble activity during SI-opposite corner stays and during freeze states of the CFC recall session, we first identified behaviour-relevant timepoints around the onset of each SI-opposite corner entry or the onset of the freezing behaviour in the CFC recall session. Pearson (for parametric data) or Spearman (for nonparametric data) correlation methods were then used to evaluate the correlation between averaged CFC recall-specific ensemble activity during the events of SI-opposite corner stay and freeze state in the CFC recall session. CFC recall-specific ensembles that show no activity in both of the two behavioural onsets were not included in this correlation analysis.

For offline awake sessions, we compared the ensemble activity rate during a 2.5-minute post-CFC awake period (30 minutes after CFC training session) to either a 2.5-minute homecage session before Neutral/SoD experience as a control session (pre-SoD or pre-Neutral) or a 2.5-minute homecage session of the post-SI awake period (post-SI, 30 minutes after SI). For offline sleep sessions, we compared ensemble activity rate during sleep within the first 2 hours post-CFC training to sleep within the first 2 hours of sleep in the homecage before the first episode of Neutral/SoD experience or sleep within the first 2 hours of post-SI. For the offline REM sleep session, mice displaying all three REM sleep sessions were only included in the analysis, as some mice did not display REM in one of the control, post-SI, or post-CFC periods.

### Co-reactivation of PCA/ICA ensemble neurons during awake and sleep

To assess offline co-reactivation of ensemble neurons associated with two distinct emotional experiences, we applied PCA/ICA methods to neural activity data from either the first day (SoD or Neutral day 1, 2.5 minutes session) or the last day (SoD or Neutral day 6, 2.5 minutes session), and another independent PCA/ICA was applied to the CFC training session subsequently. After identifying ensembles for SoD (or Neutral) and CFC training sessions, and followed by backprojection to offline sessions, we calculated co-activity between SoD (or Neutral) and CFC training-related PCA/ICA ensemble neurons, similar to previously described^20,72^.

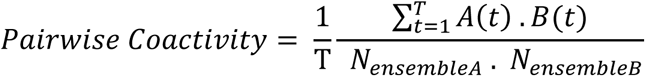

Where *A*(*t*) and *B*(*t*) are number of active neurons of ensemble A and ensemble B at timepoint *t*, *N*_*ensembleA*_and *N*_*ensembleB*_are total number of neurons per ensemble. Through these calculations, we computed pairwise co-reactivation between each ensemble A1, A2, A3…Ax from SoD/Neutral and ensemble B1, B2, B3…By from CFC training session. Results of pairwise co-reactivations of two different sets of ensembles were plotted in the matrix format (one axis showing each ensemble ID of Neutral/SoD and one axis showing the ensemble ID of the CFC training session). Co-reactivation of two independent ensembles from A and B was considered significant if neurons of ensemble Ax (SoD or Neutral session) and neurons of ensemble By (CFC training session) show co-reactivation exceeding the 99th percentile of a null distribution generated from 1000 temporally shuffled (circularly shifted) surrogate datasets. To quantify the co-activity index during post-CFC awake and sleep periods, we calculated the difference in the percentage of significant co-reactivated pairs (that is, the total number of significant pairwise co-reactivations out of the total number of all possible co-reactivations between ensembles A and B in the matrix) between post-CFC offline sessions (awake and sleep) and control offline sessions (awake and sleep) recorded before SoD or Neutral experience. For REM sleep comparison, mice displaying REM sleep in both control and post-CFC offline were only included.

### Extraction of negative emotional behaviour-related cells

For extracting behaviour-related cells, we first used the automatic behaviour analysis software EthoVision to obtain binarized behavioural timestamp data during the social interaction and CFC recall sessions. Next, we aligned the time series behavioural data with 1-second binned, z-scored neural data recorded from Ca^2+^ imaging experiments. For social avoidance-related neurons (SI avoidance), we focused on neural activity when mice were staying in the opposite corners (SI opposite corners) during encountering a novel ICR mouse in a wired-mesh cage (SI-ICR) in the open field arena. For CFC negativity-related neurons (CFC aversion), we extracted shock-related neurons, which focused on neural activity during and after electric footshock (2-second shock and 3-second post-shock), and freeze-related neurons active during the freeze state in the CFC recall session. The Kendall correlation method was used to extract single neurons that showed significantly correlated activity with each corresponding behavioural timestamp above the chance level created by circularly shifted surrogate datasets. We then used the identified neuron IDs for each behaviour-related cell type to calculate the percentage of each cell population and overlapping population out of the total number of recorded neurons per mouse.

### Functional connectivity analysis

To assess functional connectivity, we performed neuron-to-neuron pairwise correlations using the kendall correlation method with a *p*-value threshold of 0.05, calculating the percentage of significantly correlated neuronal pairs (*p* < 0.05, *r* > 0) for each condition. For behaviour-related neurons, we identified SI avoidance-related neurons (active during opposite corner stays) and CFC aversion-related neurons (shock and freeze-related neurons) using 1-second binned, z-scored neural data from Ca^2+^ imaging, aligned with binarized behavioural timestamps. We computed intercell types pairwise neuron-neuron correlations between SI avoidance-related and CFC aversion-related neurons (excluding overlapping population between SI and CFC) during the freeze and non-freeze states in the CFC recall session. A functional connectivity index was calculated by subtracting the percentage (proportion) of significantly correlated pairs during the freeze state from the percentage of correlated pairs during the non-freeze state in the CFC recall session. For connectivity index within the same SI avoidance-related cells, the percentage of significant intra-cell types pairwise correlations within SI avoidance-related cells (x and y axis with the same neuron IDs) was computed during freeze state, subtracted by the percentage during non-freeze state in the CFC recall session.

For offline analysis, we assessed functional connectivity between NREM-co-reactivated SoD-ensemble neurons (from PCA/ICA SoD ensembles significantly co-reactivated with PCA/ICA CFC training ensembles during post-CFC NREM) and CFC recall-specific ensemble neurons across pre-SoD/neutral experience and post-CFC offline sessions. Using two independent populations (NREM-co-reactivated SoD and CFC recall-specific neurons, excluding overlapping population), we calculated the percentage of significantly correlated pairs for each offline session. A functional connectivity index was computed by subtracting the percentage of significantly correlated pairs in post-CFC offline sessions from the percentage of correlated pairs in pre-SoD/neutral experience offline sessions. Data per mouse containing at least one CFC recall-specific ensemble and both offline sessions of control and post-CFC were used for analysis.

### Co-reactivation of negative behaviour-related cells during awake and sleep

To assess interaction between subtypes of negative emotional behaviour-related neurons (SI avoidance and CFC aversion-related) during post-CFC offline states, co-reactivation of these two independent sets of behaviour-related neurons was calculated, using a similar method to co-reactivations of PCA/ICA ensemble neurons. In brief, we first binarized z-scored time series neural data of the offline sessions as 1 indicating neuron active and 0 as inactive at time point t. First, the co-reactivation score was computed by multiplying the number of active neurons of SI avoidance-related and CFC aversion-related cells, normalized by the product of group sizes at each time point. These scores were averaged across all timepoints to quantify the simultaneous co-reactivation of SI avoidance and CFC aversion-related cells during the whole offline (awake or sleep) session. Finally, the co-activity index was calculated by subtracting quantified co-reactivations of post-CFC offline by quantified co-reactivations of pre-SoD (or Neutral) offline session serving as control.

### Population activity vector (PV) analysis

Neural data for each session of interest or during stimulus presentation (CS^+^, CS^−^, GS1, GS2) were averaged across timepoints to produce a single population activity vector (PV) per session, representing the mean population response during that period. For PVs representing post-SoD sleep on the last day of SoD experience (day 6), we first extracted SoD-session neurons (defined as those with at least two-fold higher mean activity during the SoD session compared to pre-SoD homecage activity on day 6) and US (shock-responsive)-related neurons from the training 2 session (Training 2) before SoD. We then pooled two types of averaged population activity during sleep: (1) neural activity during reactivation of SoD-session neurons as an ensemble-like unit and (2) neural activity during co-reactivation of SoD-session neurons with US-related neurons.

To identify and pool these reactivated and co-reactivated timepoints, we first binarized the neural data during sleep (activity was considered active if z-score ≥ 2). We then counted the number of synchronously active neurons within the SoD population (for reactivation) or the number of simultaneously active SoD neurons with US-related neurons (for co-reactivation) at each timepoint. For reactivation, candidate timepoints were those with at least four SoD neurons synchronously active, ensuring robust detection of ensemble-like unit activity while minimizing inclusion of random synchronous activations. For co-reactivation, candidate timepoints required at least two neurons from each population (SoD and US-related neurons). To determine significance, we generated 1000 surrogate datasets by circularly shifting the temporal activity of each neuron. For each neuron, we performed a permutation test on its reactivation or co-reactivation frequency, identifying significant neurons as those with real frequencies exceeding the shuffled distribution. Using only these significant neurons, we recomputed the timepoints where significant SoD neurons were reactivated as an ensemble-like unit or co-reactivated with US-related neurons.

After pooling the significant timepoints for reactivation or co-reactivation, we computed the mean population activity across these timepoints to generate the corresponding PVs (post-SoD-sleep-reactivated and post-SoD-sleep-coactivated). The PV for CS^+^ presentation after SoD (CS^+^ Recall 2) served as the reference vector. Relative cosine angles between each PV and the reference vector were computed to measure representation similarity, as previously described^72,80^. In brief, to measure representation similarity, we calculated the cosine similarity between each PV and the reference vector for each mouse, defined as:

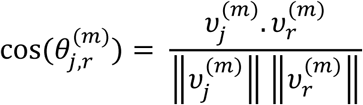

where 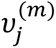 . 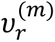 is the dot product, and 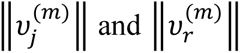 are the Euclidean norms. The angle in degrees for each mouse was computed as:

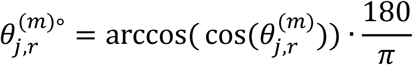

and averaged across mice. These mean angles were used to quantify the similarity of neural representations to the CS^+^ Recall 2 reference for statistical comparisons.

### Multidimensional scaling (MDS)

To visualize the global structure of representation similarity across sessions, we first computed a cosine distance matrix for each mouse’s PVs. The cosine distance between two population vectors 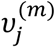 and 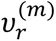 for mouse *m* is

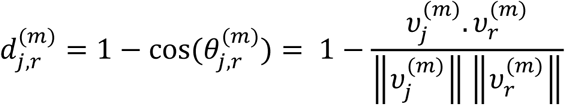

We averaged cosine distance across mice as:

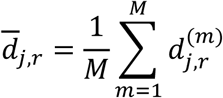

Where *M* is the number of mice.

Classical multidimensional scaling (MDS) was applied to averaged cosine distance matrix to embed PVs into a 3D space, minimizing the stress function:

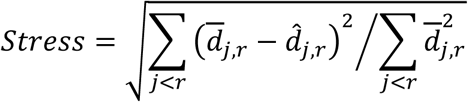

where 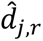 is the Euclidean distance between points *j* and *r* in the 3D embedding.

### Decoder analysis

To decode CS^+^ during Recall 2 from neural activity of sleep-active SoD subpopulations, we first filtered neural data recorded during the post-SoD recall session (Recall 2) to extract pre-identified SoD subpopulations (sleep-reactivated, sleep-co-reactivated with US, and sleep-nonreactivated), excluding neurons overlapping with CS or US-related populations. For each mouse, we trained a linear kernel support vector machine (SVM)^32,81,82^ with 10-fold cross-validation and 30 repetitions to classify CS^+^ trials (20 s CS+ and 20 s trace period) from pre-CS^+^ baseline activity at each timepoint. The baseline was defined as the mean neural response from -20 to -16 s (for 5 seconds) before CS^+^ tone onset, selected for its stable baseline response. Neural data of early trials of CS^+^ (seven trials) were used to build a decoder due to the ongoing extinction effect as the trials accumulated without US reinforcement. Decoding accuracy was calculated as the mean correct classification rate across 30 repetitions of 10-fold cross-validation for each timepoint. To compare decoding performance across mice, we extracted the peak decoding accuracy (top 20% of performance) within each 5-s block of the combined CS^+^ and trace^+^ periods and averaged these peak decoding performances per block across mice. For shuffled datasets, class labels (CS^+^ vs. pre-CS^+^ baseline) were randomized across 1000 iterations to establish chance-level performance.

To decode freeze versus non-freeze states, we trained SVMs (10-fold cross-validation with 30 repetitions) on neural data from SoD sub-populations during all CS^+^ or novel tone (GS1) presentations, including trace periods, to classify freezing versus non-freezing behaviours. To balance the number of freeze and non-freeze class labels, we applied random undersampling method (RUS) within each cross-validation fold to match the number of trials for the majority class to the minority class. Decoding accuracy was computed as the averaged classification accuracy across the 30 repeats of 10-fold CV, same as the analysis of CS^+^/pre-CS^+^ baseline procedure.

### Freezing probability around decoding timepoints

To assess freezing probability around decoding timepoints, we calculated binarized freezing scores at each second per trial for each mouse using automated behavioural analysis DeepLabCut^83^ combined with BehaviorDEPOT^84^ open-source software for freeze scoring. Freezing probability for each second was calculated by summing the number of trials with a freeze bout at each second across the 40-s CS^+^ and trace periods and dividing by the total number of trials, expressed as a percentage. For each mouse, we identified timepoints with high (top 20% of decoding performance) and low (bottom 20% of decoding performance) decoding accuracy across trials. Freezing probabilities around these timepoints were extracted within a -2 s to +2 s window relative to decoding onset and pooled across mice. To quantify changes in freezing probability, we computed the difference in AUC (Δ AUC) of freezing probabilities between post-decoding onset (0 to +2 s) and pre-decoding onset (-2 to 0 s) of all extracted timepoints.

### Histology

All experimental mice with surgeries were anesthesized and transcardially perfused with PBS followed by 4% paraformaldehyde (PFA). After brains were removed and fixed with 4% PFA for at least additional 24 hours, brains were equilibrated with 25 % sucrose solution for two days. Brains were snapfrozen with dry-ice and stored in deepfreezer (- 80°C) for future IHC. Cryosection (Leica) of frozen brains were performed with the slice thickness of 50 um and the slices were either moved to PBS solution filled cell culture plates or cryoprotectant (25% glycerol, 30% ethylene glycol, 45% PBS) solution filled plates for the long-term storage. For subsequent immunostaining, slices were washed with TBS-T solution (0.2% Triton X-100, 0.05% Tween-20) and blocked with 3% normal donkey serum (NDS) in TBS-T for 1 hour at room temperature (RT). For primary antibody treatment, 1:1000 ratio of rabbit anti-DsRed (Clonetech, #632496), 1:1000 ratio of rabbit anti-GFP (Invitrogen, #A11122) or 1:2000 ratio of alpaca anti-ALFA with Alexa647 (NanoTag, #N1502-AF647-L) were applied in blocking buffer for 24 hours at 4°C. After washing slices with PBST for three times, following secondary antibody: 1:1000 ratio of donkey anti-rabbit Alexa Fluor 546 (Molecular Probes, A10040) or 1:1000 ratio of donkey anti-rabbit Alexa Fluor 488 (Molecular Probes, A21206) along with 1:5000 ratio of DAPI in blocking solution were treated for two hours at room temperature. After washing slices with PBST for three times followed by subsequent PBS washing, slices were mounted (Kaiser’s Glycerol Gelatine or Prolong Gold Antifade Mountant) on the glass slides for fluorescent microscopy imaging. For counting the percentage of expressed ArchT-cell, total number of ArchT-expressing cells was divided by the DAPI-positive area.

### Statistical analysis

All statistical analyses were performed using GraphPad Prism (version 10) and MATLAB (2024b). The number of samples was determined based on previous studies. All data are expressed as mean ± s.e.m. All statistical tests were two-tailed. For two-sample tests, t-tests (unpaired t-test, t-test with Welch’s correction) for parametric data or Mann-Whitney U tests for nonparametric data were performed for independent comparisons, and paired t-tests for parametric data or Wilcoxon signed-rank tests for nonparametric data were performed for matched comparisons based on normality and differences in variances. For multi-sample tests, one-way ANOVA, two-way ANOVA (ordinary or repeated measures), or alternatives with multiple comparison test corrections were performed based on the normality of residuals and sphericity of data.

## Data availability

All data and resources that supported the findings of this study are available upon request.

## Code availability

All codes that supported the findings of this study are available upon request.

## Acknowledgements

We thank Kazuki Fujii, Keizo Takao and his lab/staff members of the animal facility of the University of Toyama for providing and assisting the behavioural room with automatic video tracking software; Shuhei Tsujimura and Mina Matsuo for the genotyping and IVF/maintenance of animals; the Engineering Machine Shop of the Faculty of Engineering at the University of Toyama for the fabrication of the behavioural apparatus; Takashi Takekawa (Kogakuin University of Technology and Engineering) for the automatic Ca^2+^ sorting/cell detection system HOTARU; Mostafa Reda Fayed (Inokuchi lab) for helping the preparation of AAV virus stocks and all the current/alumni laboratory members of Kaoru Inokuchi lab, Tatsuya Haga (NICT-CiNet), Taro Toyoizumi and his laboratory members (RIKEN-CBS), Takashi Kitamura (UT Southwestern), Kenta Hagihara (NIPS) and Teruhiro Okuyama (University of Tokyo) for their valuable comments and discussions. This work was supported by grants from JSPS Kakenhi (grant numbers: JP18H05213 and JP23H05476), the Core Research for Evolutional Science and Technology (CREST) program (grant numbers: JPMJCR13W1 and JPMJCR23N2) of the Japan Science and Technology Agency (JST), the Takeda Science Foundation to K.I., and the AMED PRIME (grant number: 23gm6510028h0001), JSPS KAKENHI (grant number: 20H03554 and 23H02785), the Uehara Memorial Foundation, the Naito Foundation, the Inamori Foundation, the Takeda Science Foundation, the Firstbank Of Toyama Scholarship Foundation Research Grant, the Hokuriku Bank Grant-in-Aid for Young Scientists, and the Tamura Science and Technology Foundation support to M.N.

## Author contributions

S.U. and K.I. designed the project and experiments and wrote the manuscript. S.U. and M.K.H. conducted the behavioural experiments. S.U., R.O., M.K.H., E.M., K.Y., and K.C. performed the stereotactic surgeries, perfusion and histology. S.U. performed all the Ca^2+^ imaging experiments. S.U. and M.N. performed the data analyses and wrote the MATLAB codes. M.N. generated the circuit files for the TDT system. S.U., R.O., E.M., H.A. and A.S. designed/performed the cloning and prepared the adeno-associated viral vectors. S.U. wrote the original draft of the manuscript. K.I. supervised all aspects of this work.

## Extended Figures

**Extended Data Fig. 1.**
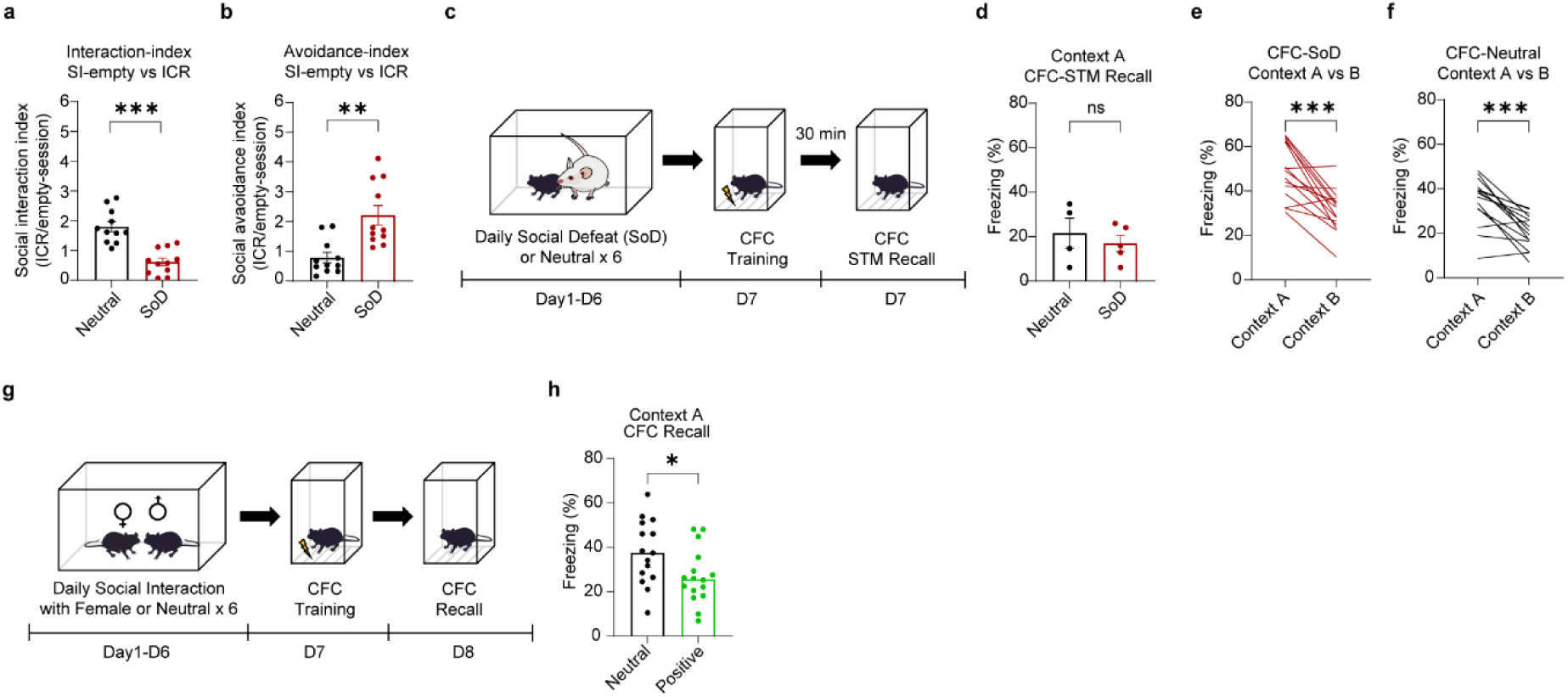
Negative emotional states induced by SoD reduced social interaction and increased avoidance indices; however, discrimination of the two contexts in CFC trials and freezing behaviour in short-term memory recall tests were comparable, whereas a positive emotional state induces lower freezing behaviour in the memory recall test after 24 hours. **a.** Social interaction index calculated by the time spent in the interaction zone when the ICR mouse was present (ICR session) divided by the time spent when the ICR mouse was absent (empty session) (Neutral, *n* = 11; SoD, *n* = 11; unpaired t test, *t* = 5.608; *P* < 0.001). **b.** Social avoidance index calculated by the time spent in opposite corners during the ICR session compared with the time spent in the empty session (Neutral, *n* = 11; SoD, *n* = 11 mice; Mann‒Whitney *U* test; *P* = 0.0019). **c.** Timeline of the SoD-short-term memory (STM) recall test paradigm. **d.** Total freezing level (%) of the neutral and SoD groups in the STM recall test (30 minutes after CFC training) (Neutral, *n* = 4; SoD, *n* = 5 mice; Unpaired t test, *t* = 0.6382; *P* = 0.5436). **e.** Total freezing percentage (%) in the recall context (context A) and novel context (context B) in the SoD group (Neutral *n* = 15; SoD *n* = 15 mice; paired t test, t = 5.135; *P* < 0.001). **f.** Total freezing percentage (%) in the recall context (context A) and novel context (context B) in the neutral group (Paired t test, *t* = 5.157, *P* < 0.001). **g.** Timeline of the positive emotional state (daily social interaction with stranger female mice for 2 hours) CFC paradigm. **h.** Total freezing percentage (%) of the neutral group and positive (positive emotional state) group in the CFC memory recall test after 24 hours (Neutral *n* = 15; SoD *n* = 16 mice; unpaired t test, *t* = 2.287; *P* = 0.0297). Data are presented as the mean ± s.e.m. Statistical significance is expressed as ns, not significant; * *P* < 0.05; ** *P* < 0.01. *** *P* < 0.001.

**Extended Data Fig. 2.**
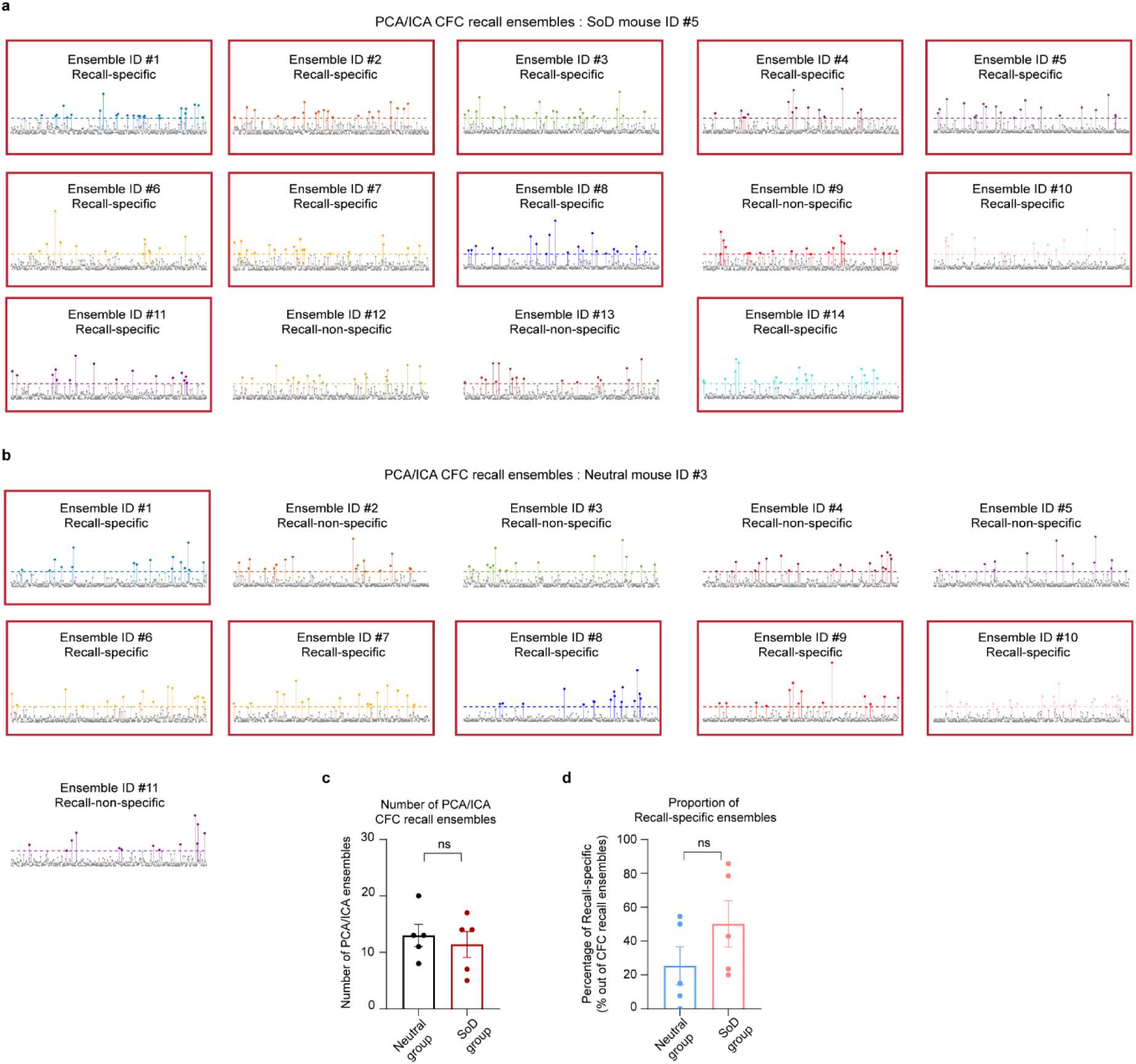
Comparison of the identified proportions of the PCA/ICA CFC recall-specific ensembles between the neutral and SoD groups. **a.** Examples showing the number of identified CFC recall-specific ensembles among all the PCA/ICA CFC recall ensembles in a single SoD mouse (mouse id #5) and **b.** Number of recall-specific ensembles in a single neutral mouse (mouse id #3). **c.** Total number of PCA/ICA CFC recall ensembles per mouse in the neutral and SoD groups (Neutral, *n* = 5; SoD, *n* = 5 mice; Unpaired t test, *t* = 0.5287; *P* = 0.6114). **d.** Proportion of identified CFC recall-specific ensembles out of all the PCA/ICA CFC recall ensembles per mouse (Unpaired t test, *t* = 1.395, *P* = 0.2006). Data are presented as the mean ± s.e.m. Statistical significance is expressed as ns, not significant.

**Extended Data Fig. 3.**
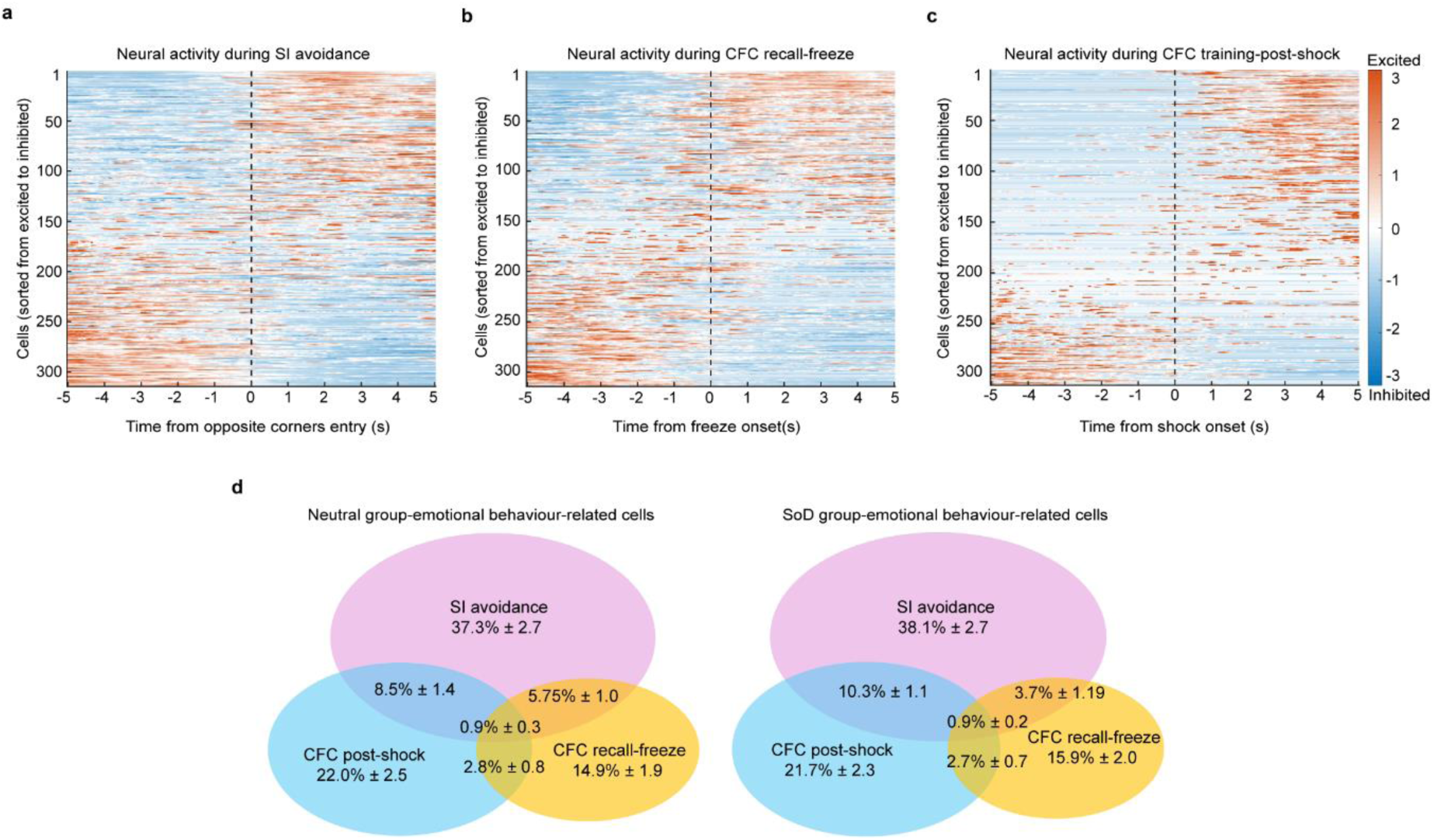
Subtypes of negative emotional behaviour-related cells, revealing no difference in the overlaps among different types of emotional behaviour-related cells between the neutral and SoD groups. **a.** Example heatmap of all recorded z scored neural activity (averaged) from a single mouse around the behavioural onset when the mouse entered the zone of the opposite corners during SI-ICR session. **b.** Example heatmap of recorded neural activity from the same mouse around freezing behavioural onsets during the CFC recall session. **c.** Example heatmap of recorded neural activity from the same mouse around the behavioural onsets receiving electric shocks during the CFC training session. **d.** No significant differences in the percentages of identified subtypes of negative emotional behaviour-related cells or overlaps of these cells between the neutral and SoD groups (unpaired t tests). Data are presented as the mean ± s.e.m.

**Extended Data Fig. 4.**
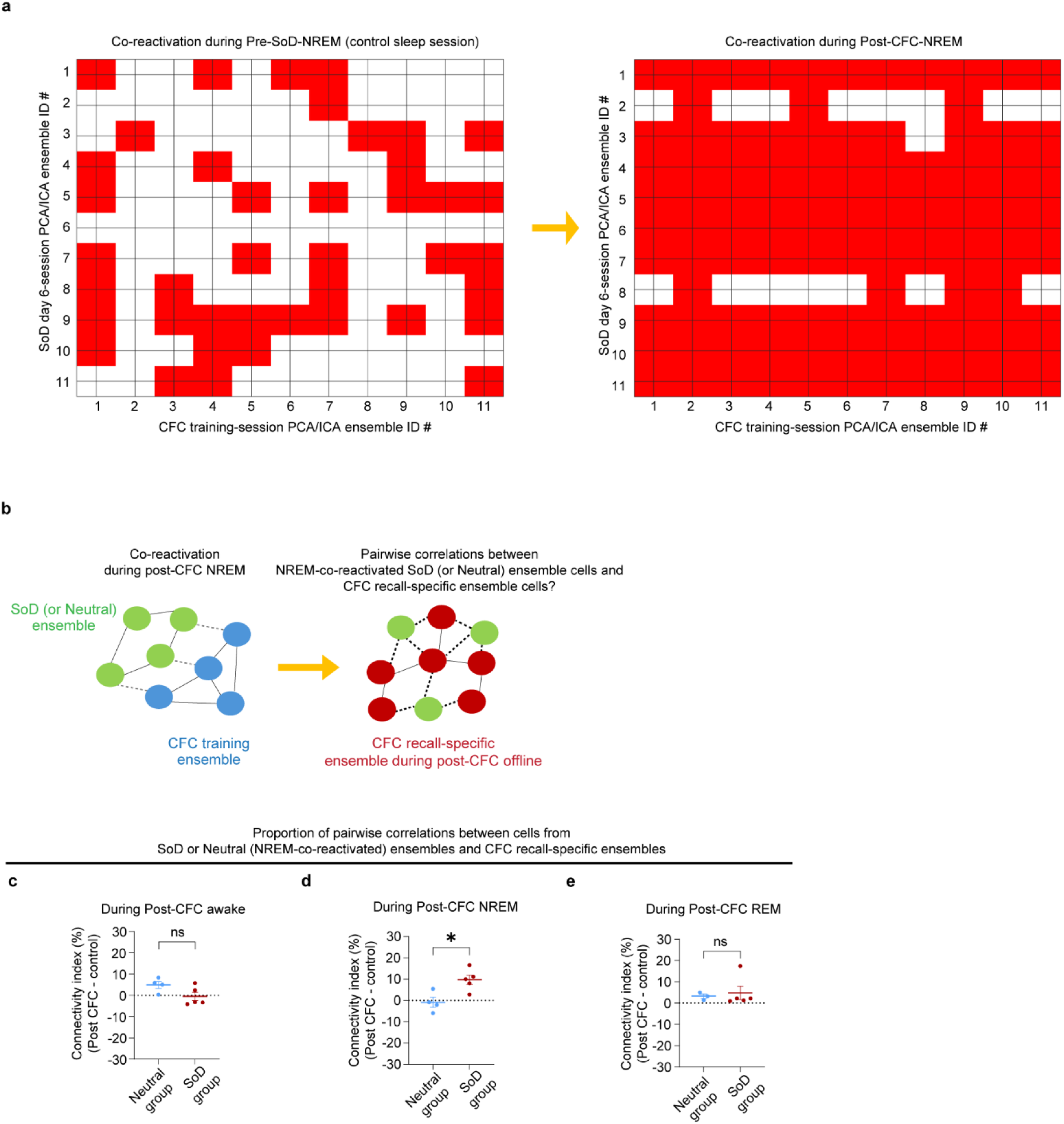
Representative heatmaps of co-reactivation of SoD session-related and CFC training session-related ensemble cells, and increased intercell functional connectivity between NREM co-reactivated SoD ensemble cells and CFC recall-specific ensemble cells during post-CFC NREM sleep. **a.** Example heatmaps of the matrix showing significant pairwise co-reactivation of PCA/ICA ensembles from the SoD day 6 session and PCA/ICA ensembles from the CFC training session during pre-SoD NREM sleep as a control (left) and post-CFC NREM sleep (right). The red square indicates significant co-reactivation between the corresponding ensemble ID of the SoD and CFC training sessions, above the chance level of the surrogate datasets. **b.** Schematic illustration of the approach to investigate how SoD (or Neutral) day 6 PCA/ICA ensemble cells (which co-reactivated with CFC training PCA/ICA ensemble neurons during post-CFC NREM sleep) interact with CFC recall-specific PCA/ICA ensemble neurons during post-CFC offline periods at the single-cell level. **c.** Percentage of significant pairwise correlations between co-reactivated SoD (or Neutral) day 6 ensemble cells (which were co-reactivated with CFC training ensembles during NREM sleep) and CFC recall-specific ensemble cells during the post-CFC awake period (Neutral *n* = 4, SoD *n* = 5 mice, Unpaired t test, *t* = 2.036, *P* = 0.0812), **d.** Same during post-CFC NREM sleep (Neutral *n* = 4, SoD *n* = 5 mice, Unpaired t test, *t* = 3.222, *P* = 0.0146) and **e.** during post-CFC REM sleep (Neutral *n* = 3, SoD *n* = 5 mice, Mann–Whitney *U* test, *P* = 0.5714). Data are presented as the mean ± s.e.m. Statistical significance is expressed as ns, not significant; * *P* < 0.05.

**Extended Data Fig. 5.**
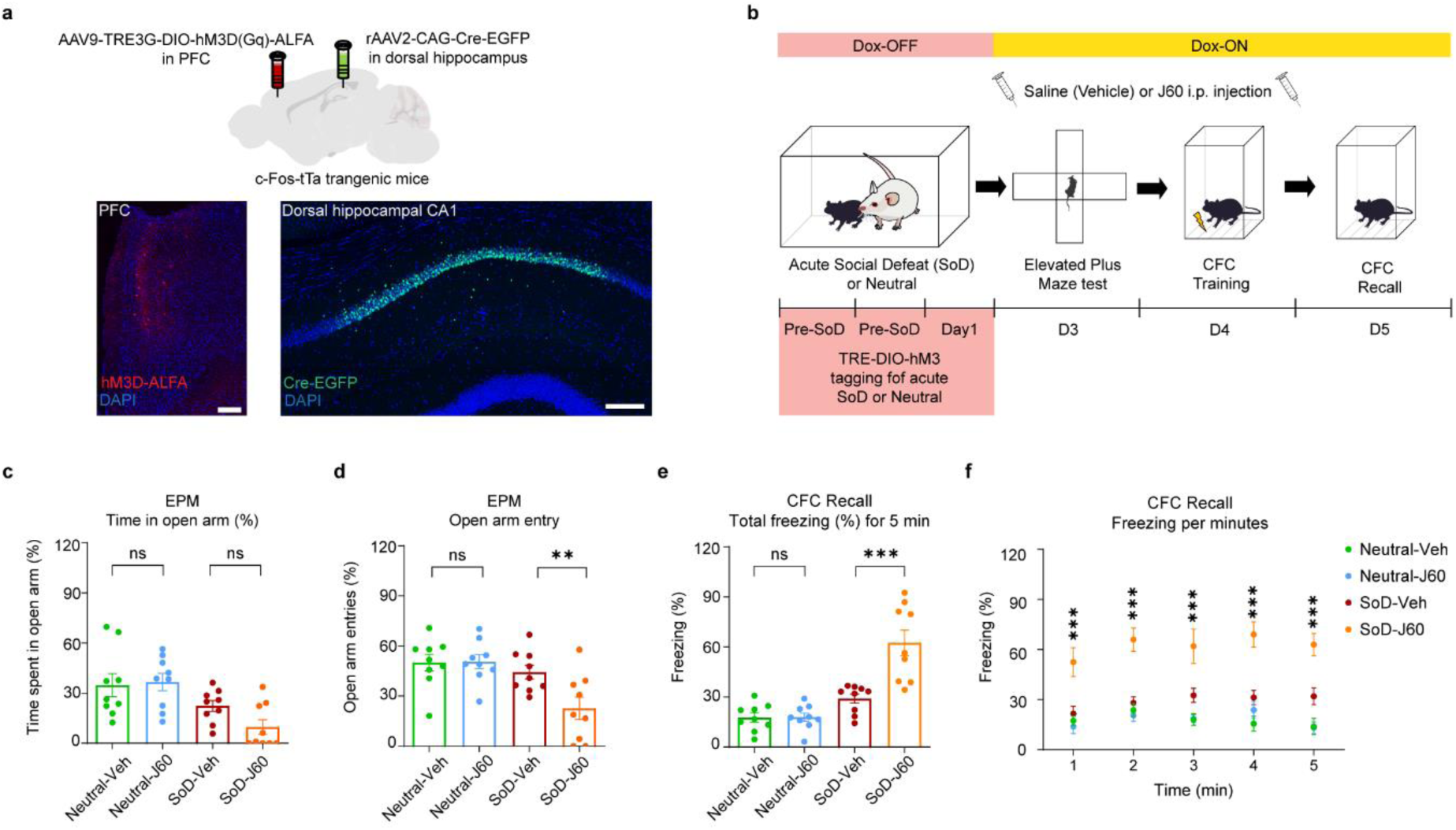
Prefrontal SoD-tagged populations projecting to the dorsal hippocampus encode anxious states and mediate negativity bias during the CFC recall test. **a.** Illustration of the virus injection strategy to retrogradely label hippocampal-projecting prefrontal cortical (PFC) populations during the SoD session and histological confirmation of the activity-dependent expression of the excitatory DREADD receptors hM3D tagged with ALFA in the PFC area (lower left) and Cre-EGFP expression in the CA1 region of the dorsal hippocampus (lower right). Scale bar, 200 µm. **b.** Timeline of the behavioural experiments to tag hippocampal-projecting PFC populations with one SoD (or Neutral) event and subsequent chemogenetic experiments in the EPM and CFC. **c.** Percentage of time spent in the open arms (EPM) in chemogenetic silencing experiments; each mouse in the neutral and SoD group was injected with either saline as the vehicle group (Veh) or the 3^rd^ generation DREADD agonist J60 (J60) as the drug group (Neutral-Veh *n* = 9; Neutral-J60 *n* =9; SoD-Veh *n* = 9; SoD-J60 *n* = 9; two-way ANOVA; interaction *P* = 0.1688; *F*(1, 32) = 1.982; Šídák’s multiple comparison test). **d.** Percentage of open arm entries in the EPM test (Two-way ANOVA, interaction *p* = 0.0353, *F* (1, 32) = 4.832, Šídák’s multiple comparisons test). **e**. Total freezing percentage (%) in the CFC recall session in the chemogenetic experiment groups (Neutral-Veh *n* = 9, Neutral-J60 *n* =9, SoD-Veh *n* = 9, SoD-J60 *n* = 9 mice; two-way ANOVA; interaction *P* < 0.001; *F*(1, 32) = 13.92; Šídák’s multiple comparisons test). **f.** Freezing (%) level per minute during the CFC recall session (statistically significant differences between the SoD-Veh and SoD-J60 groups in post hoc multiple comparison tests are shown as representative results) (two-way RM ANOVA, interaction *P* = 0.2649, *F*(12, 128) = 1.237, Dunnett’s multiple comparisons test). Data are presented as the mean ± s.e.m. Statistical significance is expressed as ns, not significant; ** *P* < 0.01; *** *P* < 0.001.

**Extended Data Fig. 6.**
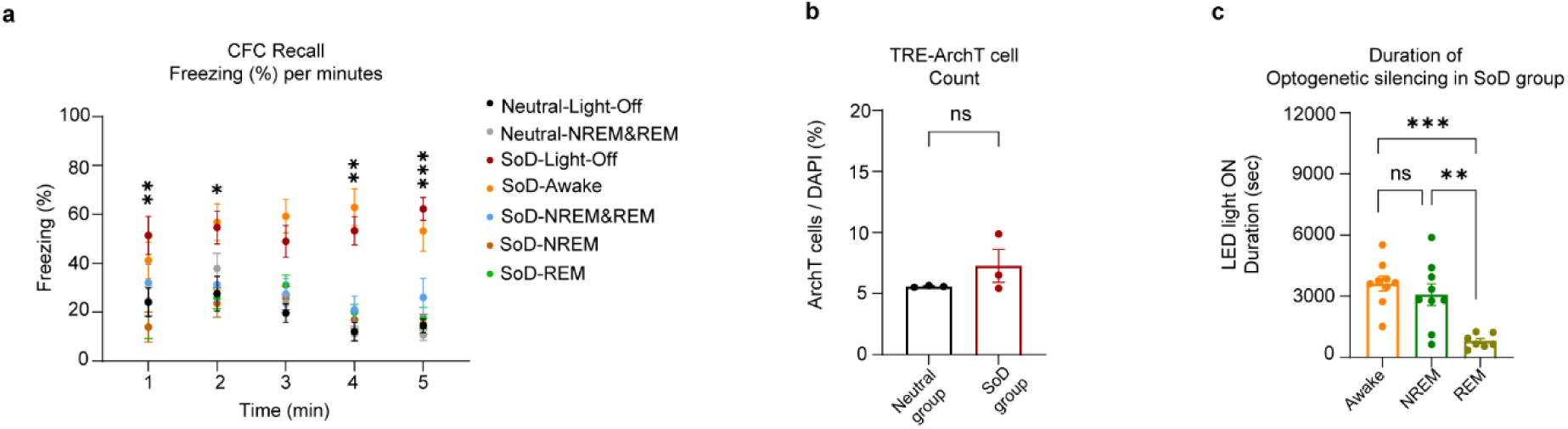
Offline sleep optogenetic silencing reduces freezing per minute in the CFC recall session among the SoD mice; the neutral and SoD groups show no significant difference in the proportion of tagged ArchT cells in the hippocampus and comparable (NREM) or shorter (REM) optogenetic silencing durations in the post-CFC sleep period. **a.** Freezing (%) level per minute during the CFC recall session (statistically significant differences between the SoD-Light-Off and SoD-REM groups in post hoc multiple comparison tests are shown as representative results) (Two-way RM ANOVA with correction, interaction *P* <0.001, *F*(24, 216) = 2.300, Dunnett’s multiple comparisons test). **b.** Percentage of ArchT-tagged cells among DAPI-positive cells in the neutral and SoD groups (Neutral, *n* = 3; SoD, *n* = 3 mice; unpaired t test with correction, *t* = 1.261; *P* = 0.3344). **c.** Duration of optogenetic silencing (LED-light ON) among the three post-CFC offline (awake, NREM and REM) manipulation groups of SoD mice (awake *n* = 9, NREM *n* = 9 mice, REM *n* = 7 mice; Brown–Forsythe ANOVA test; interaction *P* < 0.001; *F* (2.000, 15.16) = 13.59; Dunnett’s T3 multiple comparisons test). Data are presented as the mean ± s.e.m. Statistical significance is expressed as follows: ns, not significant; * *P* < 0.05; ** *P* < 0.01; *** *P* < 0.001.

**Extended Data Fig. 7.**
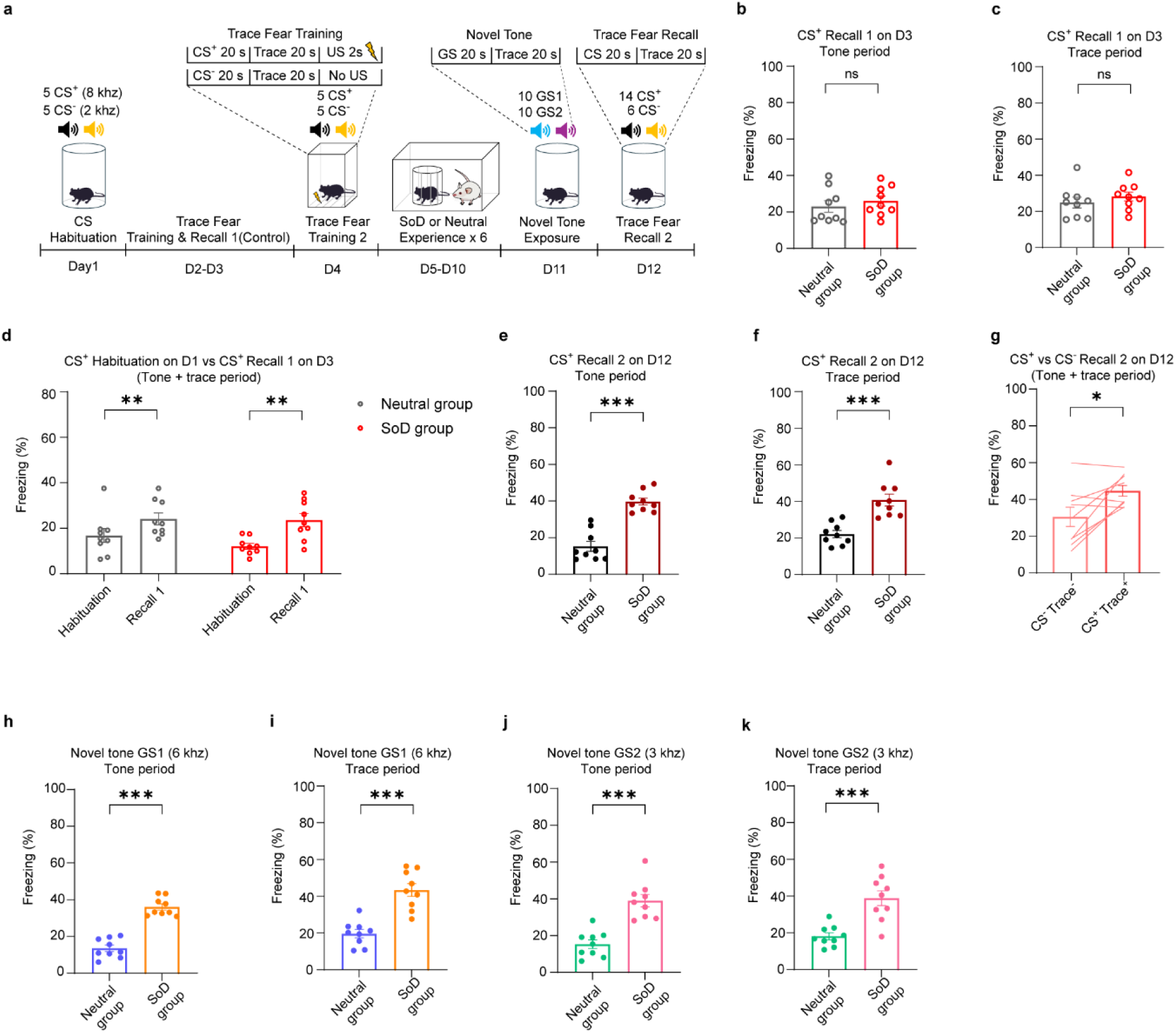
Repeated SoD experiences for six days induce negativity bias in the future recall of past trace fear conditioning (TFC) memory or ambiguous novel tones. **a.** Schematic of the timeline for the TFC training, SoD (or neutral) novel tone experiments, and TFC memory recall experiments. **b.** Total freezing (%) levels during tone presentation (each trial consisting of 20 sec) of CS^+^ (8 kHz) trials in the recall session before the neutral or SoD experiences (Recall 1 on D3) as a control session (Neutral *n* = 9, SoD *n* = 9 mice, Unpaired t test, *t* = 0.7145, *P* = 0.4852). **c.** Total freezing (%) levels during the trace period (each trial consisting of 20 sec after the termination of tone presentation) of the CS^+^ trials in recall session 1 as a control session (unpaired t test, *t* = 0.8724, *P* = 0.3959). **d.** Comparison of freezing levels during the combined CS^+^ and trace^+^ trials (each combined trial consisting of 40 sec) on the day of the CS habituation session (D1) with the freezing levels of the CS^+^ and trace^+^ trials on the day of recall session 1 (D3) for the neutral and SoD groups (Neutral-CS^+^ Habituation vs. Neutral-CS^+^ Recall, paired t test, *t* = 3.621, *p* = 0.0068; SoD-CS^+^ Habituation vs. SoD-CS^+^ Recall 1, paired t test, *t* = 4.502, *P* = 0.002). **e.** Total freezing (%) levels during tone presentation of the CS^+^ trials in the recall session 2 after the neutral or SoD experience (Recall 2 on D12) (Neutral *n* = 9; SoD *n* = 9 mice; Mann‒Whitney *U* test; *P* < 0.001). **f.** Total freezing (%) levels during the trace period of the CS^+^ trials in recall session 2 (unpaired t test, *t* = 4.872; *P* < 0.001). **g.** Comparison of the freezing levels during the combined CS^+^ and trace^+^ trials and combined CS^−^ (2 kHz) and trace^−^ trials on the day of recall session 2 in the SoD group (n = 9 mice, paired t test, *t* = 2.980, *P* = 0.0176). **h.** Total freezing (%) levels during the presentation of tone GS1 (6 kHz) in the novel tone session after the neutral or SoD experience (Neutral *n* = 9; SoD *n* = 9 mice; unpaired t test, *t* = 9.402; *P* < 0.001). **i.** Total freezing (%) levels during the trace period of the novel tone GS1 trials after the neutral or SoD experience (unpaired t test, *t* = 5.715; *P* < 0.001). **j.** Total freezing (%) levels during the presentation of tone GS2 (3 kHz) in the novel tone session after the neutral or SoD experience (unpaired t test, *t* = 5.754, *P* < 0.001). **k.** Total freezing (%) levels during the trace period of the GS2 trials of the novel tone session after the neutral or SoD experience (unpaired t test with correction, *t* = 4.639, *P* < 0.001). Data are presented as the mean ± s.e.m. Statistical significance is expressed as follows: ns, not significant; * *P* < 0.05; ** *P* < 0.01; *** *P* < 0.001.

**Extended Data Fig 8.**
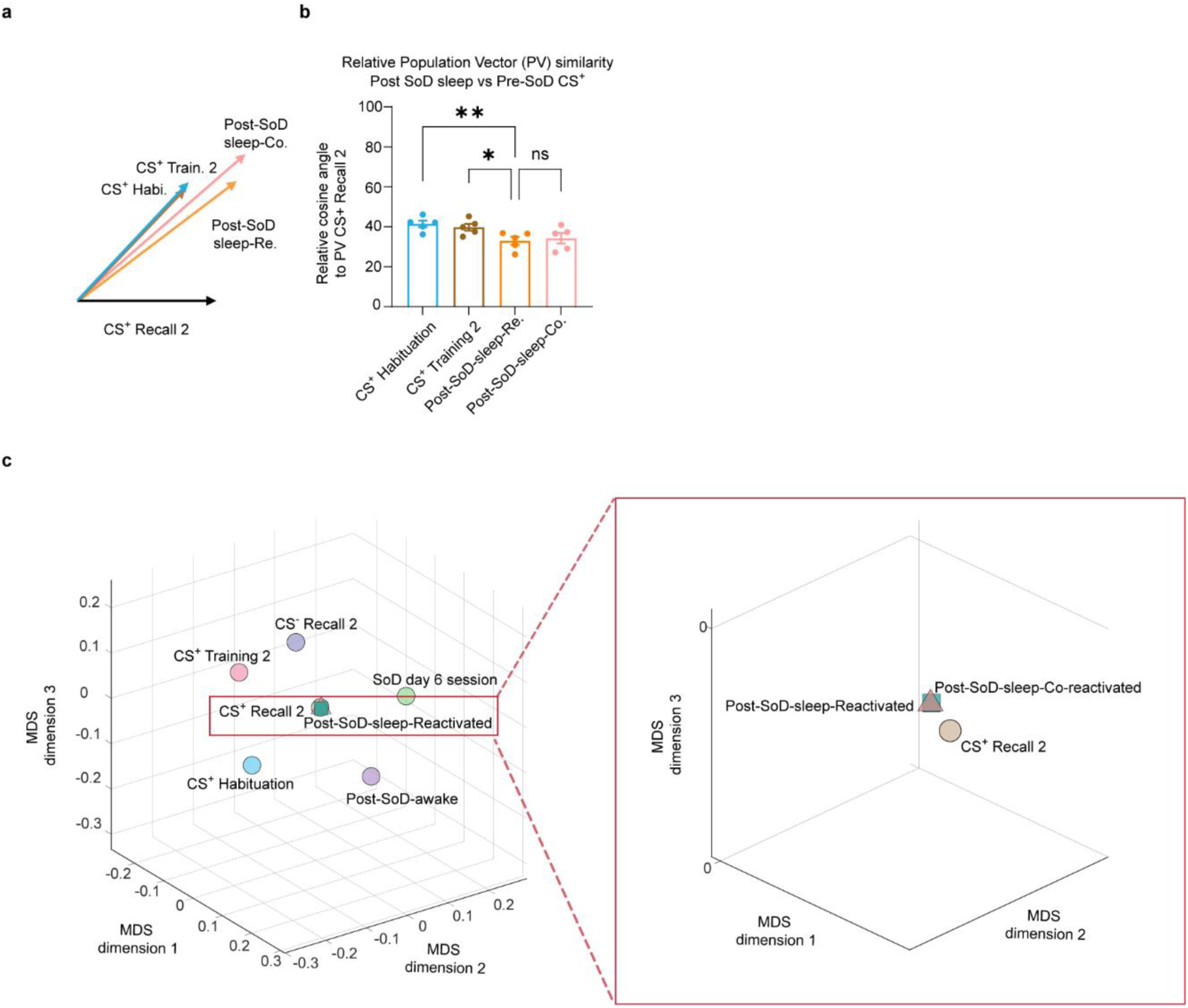
Post-SoD sleep PVs are closer to future CS^+^ PVs than are the PVs of CS^+^ sessions before the SoD experience and are clustered in the MDS space. **a**. 2D plot of the relationship between the population activity vector of the CS^+^ recall session 2 (CS^+^ Recall 2 on day 12) and the neural representation of the CS^+^ trials in the habituation (CS^+^ Habi. on day 1) and training 2 (CS^+^ Train. 2 on day 4) sessions before the SoD experience (collectively as Pre-SoD CS^+^) along with the PVs of the post-SoD sleep trials on day 10. **b.** Relative cosine angles between the reference PV (CS^+^ Recall 2) and the PVs of the pre-SoD CS^+^ and the post-SoD sleep sessions (one-way RM ANOVA, *P* = 0.0032, *F* (3, 12) = 8.101, Dunnett’s multiple comparisons test). **c.** Zoomed-in version of the MDS analysis showing that the PVs of the post-SoD sleep-reactivated, post-SoD sleep-co-reactivated and CS^+^ recall session 2 trials are clustered compared with the other PVs. Data are presented as the mean ± s.e.m. Statistical significance is expressed as ns, not significant; * *P* < 0.05; ** *P* < 0.01.

**Extended Data Fig. 9.**
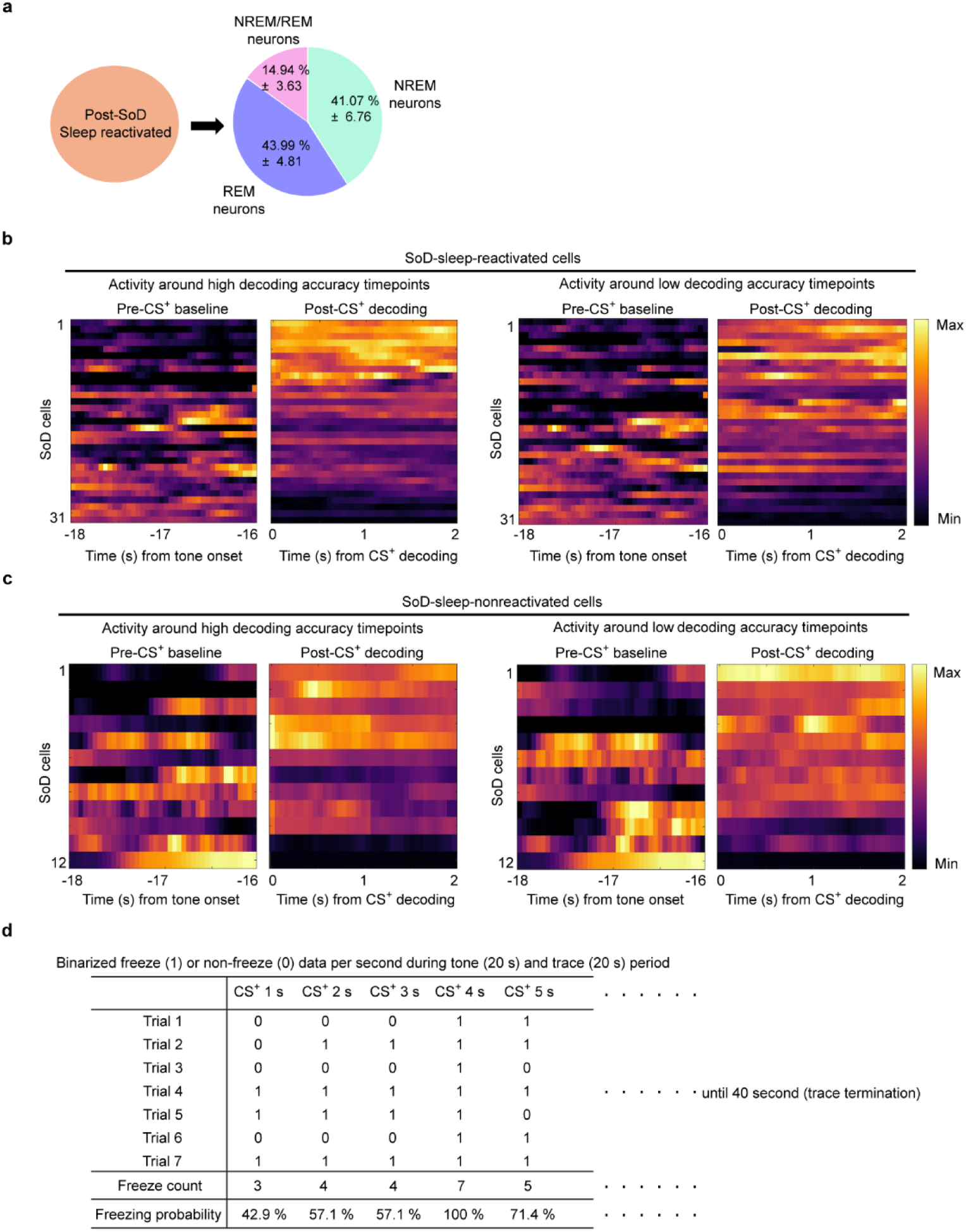
Proportion of NREM/REM-selective cells in post-SoD sleep-reactivated populations, representative neural activities around timepoints of high/low decoding accuracy in the CS^+^/baseline trials and calculation of the freezing probability of the mice during the CS^+^ and trace^+^ trials of recall session 2. **a.** Proportion of NREM-responsive and REM-responsive cells among post-SoD (day 6) sleep-reactivated cells. **b.** Representative example heatmaps of trial-averaged neural activity of SoD sleep-reactivated cells during the last 2 seconds of the pre-CS baseline response (-20 to -16 seconds from the onset of the CS tone) and 2 seconds after the CS decoding timepoints at which the decoding accuracy was relatively high (top 20% decoding performance, left) or relatively low (bottom 20% decoding performance, right). **c.** Representative example heatmaps of the trial-averaged neural activity of SoD sleep-nonreactivated cells during the last 2 seconds of the pre-CS baseline response and 2 seconds after the CS decoding timepoints at which the decoding accuracy was relatively high (left) or low (right). **d.** Example illustration of calculating the freezing probability of individual mice at each second of the tone and trace periods of the CS^+^ and trace^+^ trials in the post-SoD recall period (Recall 2).

**Extended Fig 10.**
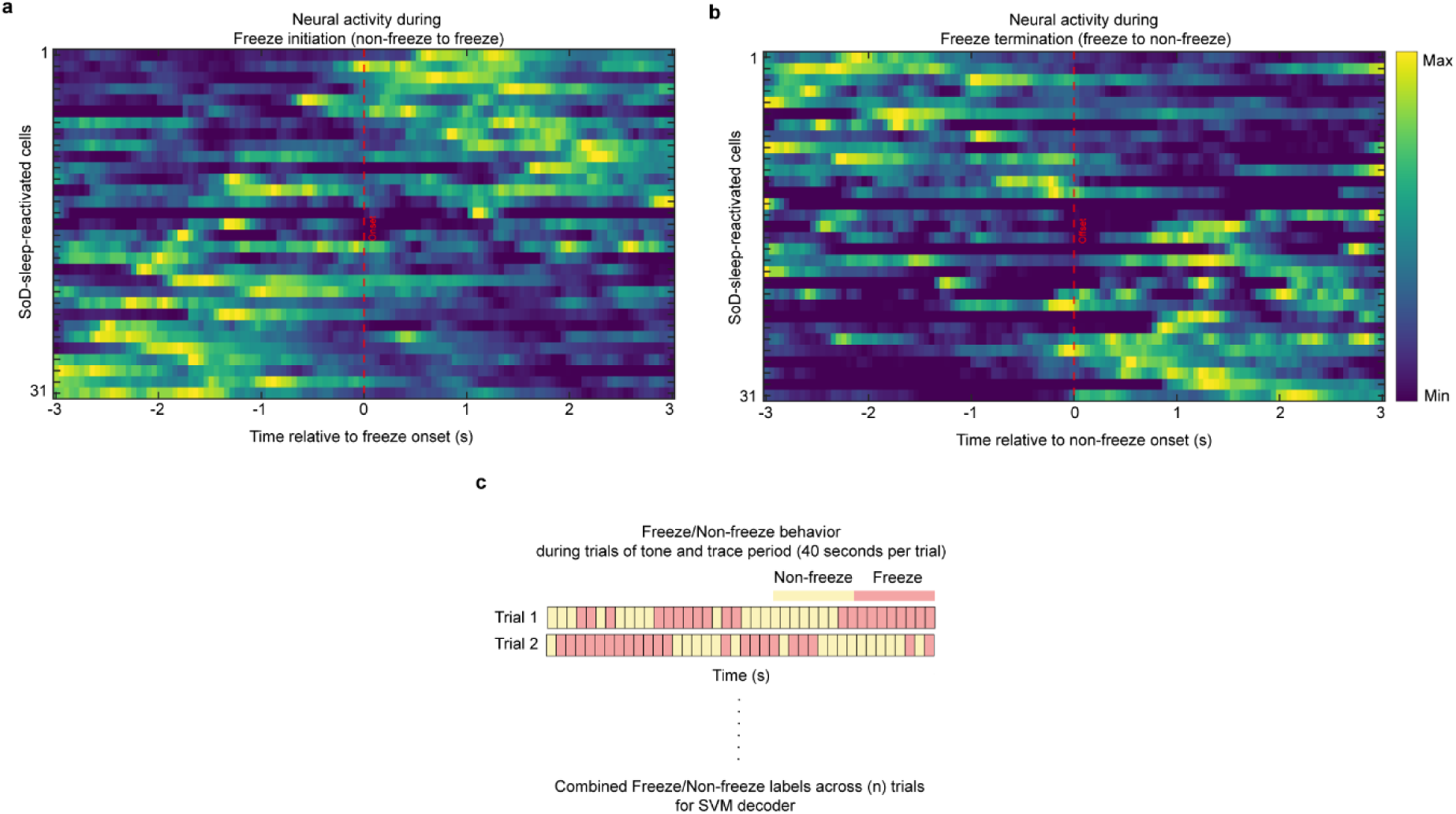
Neural activity of SoD sleep-active neurons during freeze and non-freeze states and a graphical illustration of the construction of the SVM freeze/non-freeze decoders during the combined CS^+^ and trace^+^ trials. **a.** Representative heatmap of the trial-averaged neural activity of the SoD sleep-reactivated population during non-freeze to freeze transition periods. **b.** Example heatmap of the neural activity of the SoD sleep-reactivated population during freeze to non-freeze transition periods. **c.** Schematic of the process of extracting behavioural timestamps of freeze/non-freeze states across all CS^+^ and trace^+^ trials for building the trial labels of the SVM freeze/non-freeze decoders.

